# dirCLIP profiles variant-specific RNA-protein interactions via nanopore long-read sequencing

**DOI:** 10.64898/2026.01.22.701028

**Authors:** Mohamed Bahrudeen, Mauro Scaravilli, Zileena Zahir, Katrina Woodward, Timo Ahola, Madara Ratnadiwakara, Jan Kaslin, Fionna Loughlin, Nikolay E. Shirokikh, Minna-Liisa Änkö

## Abstract

RNA binding proteins (RBPs) control gene expression through their activities in essentially all processing steps of coding and noncoding RNAs and are dysregulated in disease. With >95% of human multi-exon genes undergoing alternative splicing, the inability to assign RBP sites to distinct RNA variants fundamentally limits our understanding of RNA regulation. We present dirCLIP, a method combining UV crosslinking and immunoprecipitation (CLIP) with amplification-free direct nanopore long-read sequencing preserving full-length transcripts. dirCLIP employs two complementary strategies: direct RNA sequencing detecting amino acid adduct-induced perturbations in current signals and direct cDNA sequencing capturing binding sites as mutations. Benchmarking dirCLIP with SRSF3 demonstrated >75% concordance with short-read-based data, revealed isoform-selective RNA binding and enabled detection of co-occurring binding sites in single RNA molecules. Application to the noncanonical RBP HMGA1 uncovered new AT-hook mediated RNA variant-specific interactions. dirCLIP fundamentally transforms our ability to interrogate RNA regulation of distinct isoforms and transcript variants with direct implications to disease mechanisms and development of RNA therapeutics.

---

RNA binding proteins (RBPs) orchestrate every aspect of RNA metabolism, from transcription and splicing to RNA localization, translation, and decay ^1^. Understanding how RBPs selectively interact with specific RNA isoforms and transcript variants is critical for deciphering both normal physiology and disease mechanisms, with dysregulated RNA processing implicated in neuropsychiatric diseases, cancer and development ^2–6^. Our ability to map RBP interactions remains fundamentally limited by methodological constraints. Current crosslinking and immunoprecipitation (CLIP) methods for mapping cellular RBP interaction sites in RNA are based on short-read sequencing that fail to capture RNA-protein interactions in the native transcript context ^7–13^. They utilize the ability of short wave ultraviolet (UV) radiation to induce covalent bond formation between closely (within a few ångströms) placed aromatic rings found in RBPs and their target RNAs ^14^. The RBP interaction sites within RNAs can be detected by short-read RNA sequencing as read termination sites caused by the amino acid adduct left at the crosslink site after proteinase K digestion ^7,10–12^. However, conventional CLIP methods cannot determine RBP sites in full-length RNAs, they introduce PCR biases often leading to overrepresentation of the most abundant RNA species and easily amplified fragments and cannot attribute binding sites to distinct RNA variants ^15–17^. Information about binding site combinations within individual transcripts is lost, limiting our understanding of cooperative or competitive RBP interactions that govern native full-length RNAs.

The emergence of nanopore direct long-read sequencing technology presents an unprecedented opportunity to overcome these limitations ^18–20^. Here we establish dirCLIP (<u>d</u>irect identification of <u>R</u>NA binding sites by <u>C</u>ross<u>L</u>inking and Immuno<u>P</u>recipitation) utilizing nanopore long-read sequencing and enabling the mapping of RNA-protein interactions in full-length RNAs including distinct RNA variants. Our dual approach employs either direct RNA sequencing (dRNA-seq) detecting crosslink sites through signal perturbations and premature termination events, or amplification-free direct cDNA sequencing (dcDNA-seq) with strand-switching reverse transcription capturing mutation signatures across entire transcripts. We rigorously benchmarked dirCLIP in multipotent P19 cells using the well-characterized RBP Serine/arginine-rich splicing factor 3 (SRSF3) having extensive functionally validated binding sites ^21–23^. Our results reveal that RBP interactions can be isoform-specific, with binding sites restricted to single splice variants - a finding invisible to conventional short-read approaches. Furthermore, assigning RBP sites to individual long-reads revealed co-occurrence of multiple binding sites within the same RNA molecule. To demonstrate broader applicability, we investigated the RNA interactions of HMGA1, a non-canonical RBP with emerging roles in RNA regulation ^24–26^, and discovered variant-specific RNA binding within coding and noncoding RNAs. In conclusion, dirCLIP provides the first method capable of interrogating the full complexity of RNA-protein regulatory networks in their native transcript context and opens new avenues for understanding post-transcriptional gene regulation and developing isoform-specific therapeutics.

## RESULTS

To develop a method for mapping RBP binding sites in full-length RNAs, we envisioned two approaches. RBP-RNA crosslink sites could be detected either by dRNA-seq as RNA modifications within the sequencing reads or by dcDNA-seq using mutation profiling (**Figure 1A**). To examine the validity of these approaches, we employed the well-characterized P19 SRSF3-BAC cell line expressing EGFP-tagged SRSF3 with extensive short-read iCLIP reference data available, including many functionally validated RNA binding sites ^21–23^. Following *in vivo* UV crosslinking, GFP-Trap magnetic beads were used to immunoprecipitate SRSF3-bound RNAs in modified iCLIP conditions omitting RNase treatment (**Supplementary Figure 1A**) ^27^. A thorough Proteinase K peptide digestion was employed to trim the protein crosslinks to short amino acid adducts compatible with dRNA-seq and reverse transcriptase (RT) read-though followed by mutation profiling. This material, together with untreated (-UV) and UV crosslinked (+UV) total transcriptome controls, was used in the dRNA- and dcDNA-seq library preparation.

**Figure 1.**
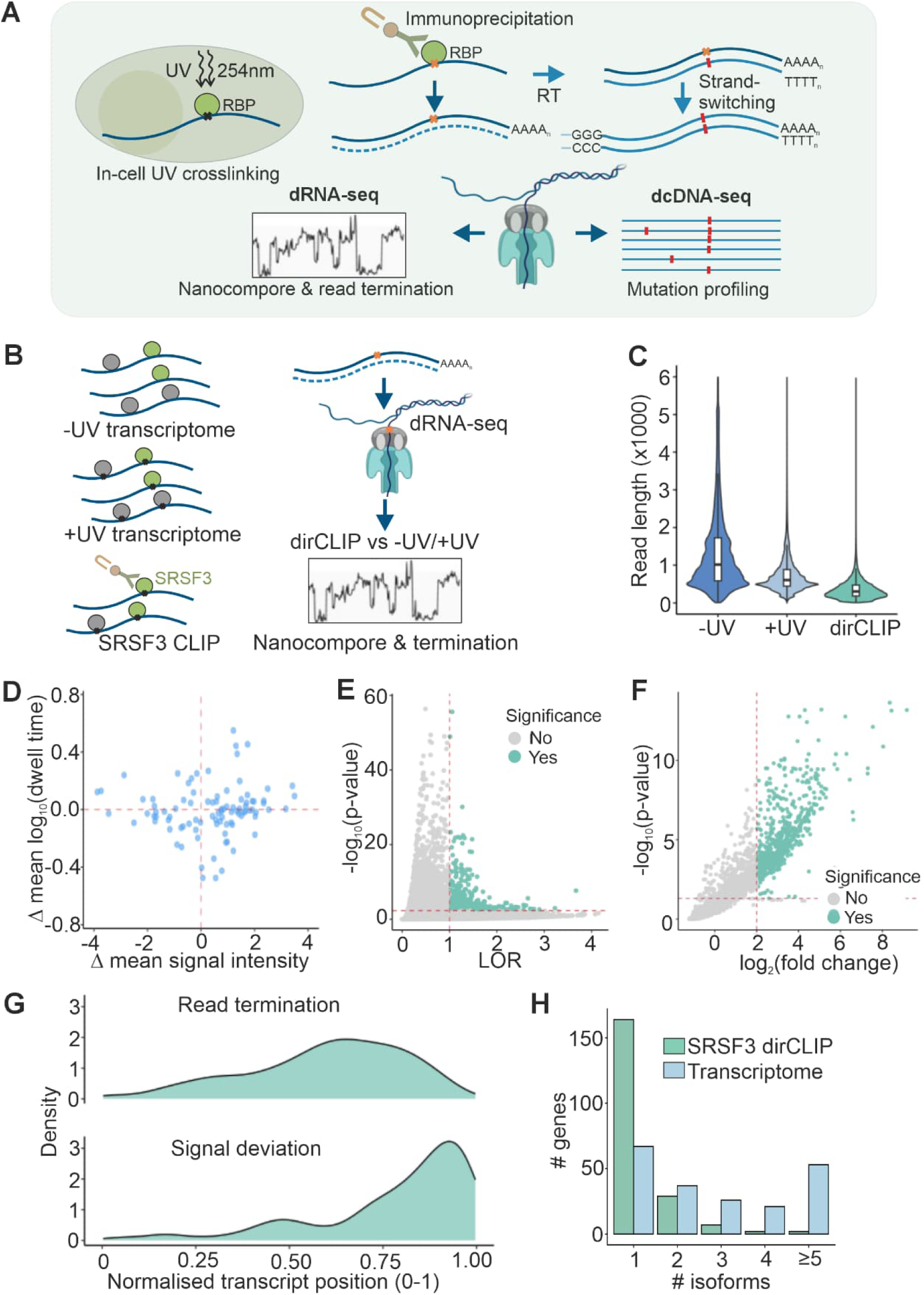
dirCLIP, <u>d</u>irect identification of <u>R</u>NA binding sites by <u>C</u>ross<u>L</u>inking and <u>I</u>mmuno<u>P</u>recipitation, for mapping of RNA-protein interactions in full-length RNAs. (**A**) Overview of the two complementary approaches utilizing nanopore direct RNA-sequencing (dRNA-seq) and cDNA sequencing (dcDNA-seq). (**B**) Overview of the dRNA-seq -based dirCLIP. (**C**) Read length distribution of untreated (-UV) and UV-treated (+UV) dRNA-seq control transcriptomes and SRSF3 dirCLIP sample. Reads <20 nucleotides or >99th percentile were excluded. (**D**) Scatter plot showing the difference in mean signal intensity (Δ mean signal intensity) versus the difference in log_10_-transformed dwell time (Δ log_10_ mean dwell time) across hundred randomly selected sites significant by the Nanocompore analysis (+UV/SRSF3). (**E**) Significant differences in signal intensity and dwell time (signal deviation) identified using Nanocompore between SRSF3 dirCLIP and UV-treated transcriptome (|LOR|>1, p-value <0.005). (**F**) Significant changes in read end termination rates between SRSF3 dirCLIP and UV-treated transcriptome detected using a Poisson GLM model (log₂FC >2, p-value <0.05). (**G**) Density distribution of significant sites identified by read termination and signal deviation methods across normalized transcript coordinates. (H) Comparison of the number of isoforms per gene with significant SRSF3 binding sites derived from SRSF3 dirCLIP versus the number of expressed isoforms (TPM > 0.5) for the same genes in the UV-treated transcriptome sample.

### Nanopore direct RNA sequencing-based detection of RNA-protein interactions

We hypothesized that covalently attached amino acid adducts would create distinctive perturbations of the nanopore read signatures detectable as altered current signals and physical blockages of RNA traversing through the nanopores. To collect the dRNA signals, the full-length RNAs recovered from the magnetic beads and total transcriptome controls (-/+ UV) were used to prepare dRNA-seq libraries with the native 3’ poly(A) (**Figure 1B**). Global analysis revealed a pronounced reduction in the read length following UV crosslinking, with SRSF3 immunoprecipitated samples showing further shortening of the reads compared to the UV-treated transcriptome controls (**Figure 1C**). These data demonstrated the successful capture of crosslinked RNA-protein complexes in our libraries but also suggested the crosslinked RNAs may result in an incomplete chain translocation through the nanopores.

Further analysis of the mapped reads revealed two complementary signatures in the dRNA-seq datasets. First, protein adducts induced characteristic changes in ionic current patterns, manifesting as alterations in both signal intensity and dwell time (**Figure 1D-E**). Second, bulky protein-RNA crosslinks caused premature read termination as the modified RNA-protein complexes failed to fully translocate through the nanopore constriction (**Figure 1F**). For the signal-based detection, we employed Nanocompore that uses a Gaussian Mixture Model (GMM) to identify positions where crosslinking significantly alters the biophysical properties of RNA translocation ^28^. The algorithm detected subtle but consistent changes: while signal intensity differences were modest, dwell times showed marked increases in SRSF3 dirCLIP samples, suggesting protein adducts slowed RNA movement through the pore (**Supplementary Figure 1B-C**). The GMM’s multivariate approach captured sites exhibiting either large changes in a single parameter or coordinated subtle shifts across multiple features (**Figure 1D**). A combination of log odds ratio (LOR) and p-value cut-offs were applied to identify significant signal alterations indicative of SRSF3 binding (**Figure 1E**).

To systematically map termination events, we developed a Poisson generalized linear model that accounts for the natural 3’ bias in nanopore sequencing while identifying positions with statistically elevated termination rates. While UV-treated transcriptome samples showed dispersed low-level termination events, reflecting known low efficiency (1-5%) of UV crosslinking ^15^, the enriched SRSF3-bound crosslinked RNAs exhibited pronounced termination enrichment in the 3’ regions, consistent with the known binding preferences of SRSF3 (**Figure 1F, Supplementary Figure 1D**).

While providing a conceptual proof-of-principle on native RNA, both detection methods were limited in their discovery capacity towards the 5’ ends of transcripts (**Figure 1G**). This gradient reflects the compound effect of natural 3’ bias in the dRNA-seq and crosslink-induced premature termination preventing the sequencing of the upstream regions. Consolidation of spatially clustered significant sites around individual binding events (**Supplementary Figure 1E**), collapsing sites within ±5 nucleotides, yielded 359 sites through signal deviation and 379 through termination analysis (total 742 sites), with only 4 sites identified by both methods (**Supplementary Table 1**). This minimal overlap suggests these approaches capture distinct aspects of protein-RNA crosslinking: signal changes likely reflect smaller adducts that permit translocation, while termination events indicate more substantial modifications that block pore passage, but are not recorded as signal deviations as they do not translocate through the pore. While using untreated (-UV) transcriptome as the control yielded more sites (**Supplementary Figure 1F-H**), motif analysis suggested that the UV-treated control improved specificity (**Supplementary Figure 1I-J**). Unlike short-read RNA sequencing, long-read sequencing is particularly adept at resolving complete mRNA isoforms that can be supported by individual reads. Strikingly, SRSF3 dRNA-seq dirCLIP sites were mostly found in single isoforms, suggesting isoform specific binding (**Figure 1H, Supplementary Figure 1K**).

Our findings provide the first example of directly observed RNA-protein crosslinks at the full-length, single-molecule RNA level with high-throughput capability and demonstrate the potential of dRNA-seq in detecting protein-RNA interactions in distinct RNA variants. These interactions detected directly in the native RNA provide complementary information through distinct biophysical signatures. However, the observed and persistent 5’ bias prompted the development of amplification-free cDNA approach to achieve comprehensive transcriptome-wide coverage.

### Direct cDNA sequencing and mutation profiling-based discovery of RNA binding sites in full-length RNAs and distinct isoforms

To enhance the detection of RBP sites across the full transcript length, we next examined the utilization of nanopore dcDNA-seq for RBP site detection. Some reverse transcriptase (RT) enzymes can read through nucleotides with adducts, introducing mutations or deletions in the copy strand at the positions corresponding to the modified nucleotide sites of the original RNA strand ^29–33^. We applied the Moloney Murine Leukemia Virus (MMLV)-based RT enzyme Maxima H Minus that is highly processive with efficient read-through across modified nucleotides and can produce double-stranded cDNA from full-length RNA molecules *via* strand-switching ^32,33^. In the dcDNA-protocol, SRSF3-bound crosslinked long RNAs were immunoprecipitated and dcDNA-seq libraries prepared from the recovered RNAs (**Figure 2A**). As controls, we sequenced P19 transcriptomes with and without UV treatment (+UV and -UV, respectively). In line with our hypotheses, the UV-crosslinked dcDNA libraries yielded markedly longer reads compared to the corresponding dRNA-seq samples (**Figure 2B-C**). Furthermore, the UV-treatment increased mutation rate with the highest rates in the dirCLIP samples (**Figure 2D**), suggesting that UV crosslinking induced mutations could identify specific RBP sites in the long RNAs as hypothesized. Deletions constituted a large proportion of the observed UV-dependent nucleotide sequence alterations (**Figure 2D**). The use of manganese (MnCl_2_) buffer conditions in the reverse transcription reaction increased the mutation rate across samples, particularly background mutations in the untreated (-UV) transcriptome control (**Supplementary Figure 2A**). Since the read length and quality (estimated by N50 values) were reduced in the MnCl_2_ conditions (**Supplementary Figure 2B**), we concluded that the standard magnesium (MgCl_2_) buffer conditions were better suited for RBP site mapping. Next, mutation profiling was conducted to identify putative SRSF3 binding sites by assessing the statistical enrichment of mutations and deletions at each position in dirCLIP compared to transcriptome samples (FDR<0.05).

**Figure 2.**
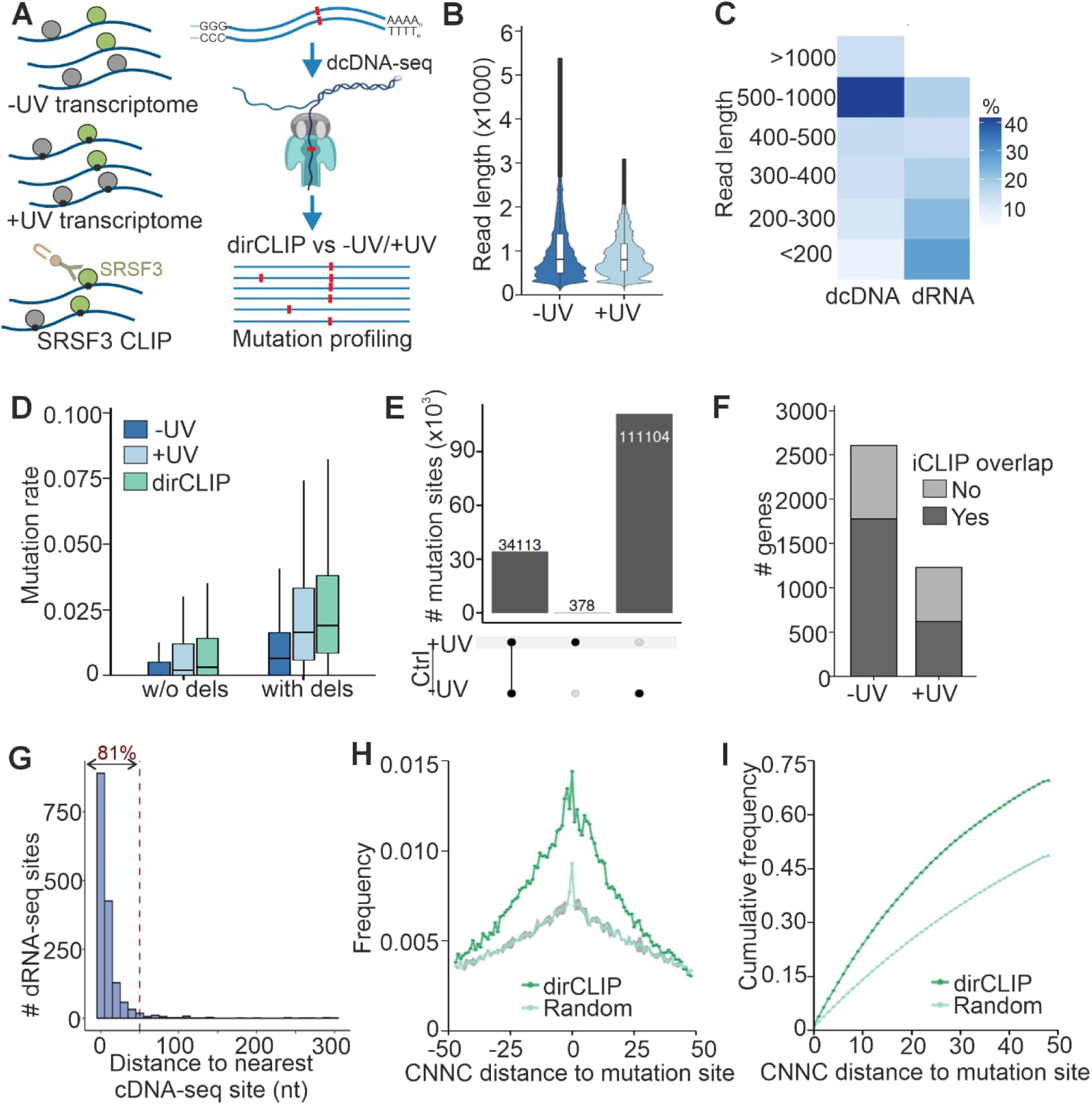
Direct cDNA sequencing-based detection of SRSF3 RNA binding protein sites. (**A**) Overview of the direct cDNA-sequencing (dcDNA-seq) -based dirCLIP. (**B**) Read length distribution of untreated (-UV) and UV-treated (+UV) dcDNA-seq control transcriptomes. Primary alignment reads >20 nucleotides or <99th percentile were used. (**C**) Read length distribution of SRSF3 dirCLIP samples sequenced using dRNA-seq and dcDNA-seq. Reads <20 nucleotides were excluded. (**D**) dcDNA-seq mutation rate per nucleotide with and without (w/o) deletions (dels) for untreated transcriptome (-UV), UV-treated transcriptome (+UV), and SRSF3 dirCLIP samples. Nucleotides with a coverage ≥50 were included. (**E**) Significant SRSF3 dcDNA-seq dirCLIP sites identified using untreated transcriptome (-UV) or UV-treated transcriptome (+UV) controls. Mutation rates were calculated including deletions and significant sites defined using a nucleotide coverage ≥50 and a mutation rate difference ≥0.03. (**F**) Overlap between genes harboring SRSF3 dcDNA-seq dirCLIP sites and iCLIP peaks when untreated transcriptome (-UV) or UV-treated transcriptome (+UV) was used as the control. Deletions were included in the calculation of mutation rates and significant sites defined using a nucleotide coverage ≥50 and a mutation rate difference ≥0.03. (**G**) Distance of significant SRSF3 dRNA-seq dirCLIP sites to the nearest dcDNA-seq dirCLIP site. (**H**) Distance between SRSF3 dcDNA-seq dirCLIP sites and CNNC motifs. As a control, sequences within ±50nt window around each dirCLIP site were randomly shuffled (random). The gray shaded area indicates the standard error of the mean from 100 rounds of random shuffling. Significant dirCLIP sites were identified using the untreated transcriptome as a control with a nucleotide coverage ≥50 and a mutation rate difference ≥0.03 including deletions in the mutation rate calculation. (**I**) Cumulative frequency distribution of SRSF3 dcDNA-seq dirCLIP site distance to the closest CNNC motif as in (H).

To evaluate the contribution of background mutations introduced by sequencing errors, we compared two independent sets of untreated P19 transcriptomes (**Supplementary Figure 2C**). We further examined the effect of read coverage and mutation difference thresholds by benchmarking our SRSF3 binding sites detected with mutation profiling against published SRSF3 iCLIP data ^12,22,27^. As expected for specific sites, increasing the read coverage threshold or mutation difference threshold reduced the number of dirCLIP sites detected. While the overlap between short-read iCLIP sites and background sites rapidly declined with more stringent thresholds, the overlap between iCLIP and dcDNA-seq dirCLIP sites remained unaffected, suggesting the putative binding sites assigned based on the mutation profiling reflected specific SRSF3-RNA interactions (**Supplementary Figure 2C**). A read coverage above fifty reads per nucleotide was sufficient to yield specific dirCLIP sites and 3% mutation difference threshold could distinguish the specific sites from background sites.

Having defined thresholds to call RBP sites, we next mapped SRSF3 binding sites using either untreated (-UV) or UV-treated (+UV) transcriptome as the control in the mutation profiling (**Figure 2E, Supplementary Table 2**). We also merged closely positioned sites into regions similar to the dRNA-seq and the published iCLIP approaches (**Supplementary Figure 2D**) ^12,27^. Applying the untreated transcriptome control enhanced the detection of specific RBP sites as demonstrated by the increased overlap with iCLIP sites (**Figure 2F**). Analysis of global positional patterns revealed that most SRSF3 dcDNA-seq dirCLIP sites were located in exons consistent with previous iCLIP data and known SRSF3 functions in facilitating processes such as exon inclusion and mRNA export (**Supplementary Figure 2E**) ^21–23,34^. The comparison of the two complementary dirCLIP strategies highlighted the specificity of the method as 81% of the dirCLIP sites detected by dRNA-seq had a corresponding dcDNA-seq site within 50nt distance (**Figure 2G**).

To further evaluate specificity, we defined SRSF3 top CNNC binding motifs based on the iCLIP peaks ^12,22,27^ and then mapped the distance of CNNC motifs to iCLIP or dirCLIP sites. Over 75% of iCLIP peaks had a CNNC motif within fifty nucleotides of the peak center (**Supplementary Figure 2F-G**). A similar proportion of the dcDNA-seq based dirCLIP sites had a CNNC motif within the same fifty nucleotide window both when using the untreated and UV-treated transcriptome as the control (**Figure 2H-I, Supplementary Figure 2H-I**), further suggesting that the mutation sites indeed reflected SRSF3 binding sites. Although deletions are considered a common type of sequencing errors in nanopore reads ^31^, we included them in the analysis because they constituted a significant proportion of sequence alterations caused by the amino acid adducts and excluding deletions in mutation profiling reduced the ability to distinguish specific sites from random sites (**Figure 2D**, **Supplementary Figure 2J-K**), suggesting that RT enzymes frequently skip nucleotides when encountering amino acid adducts.

To investigate if dcDNA-seq dirCLIP enabled the analysis of binding site co-occurrence in single RNA molecules and the detection of RBP sites in distinct mRNA isoforms, we conducted the mutation profiling by uniquely assigning the reads to transcript isoforms. Filtering of mutation sites in individual reads to retain only those overlapping with significant dirCLIP sites revealed over 50% of the reads having more than two dirCLIP sites, suggesting multiple SRSF3 bound to single RNA molecules (**Figure 3A**). Supporting the dRNA-seq dirCLIP data, SRSF3 dcDNA-seq dirCLIP sites were mostly found in single isoforms (**Figure 3B**), suggesting that SRSF3-RNA interactions may indeed be limited to selected isoforms in cells. This isoform-selective binding has remained invisible to short-read CLIP methods that largely cannot assign reads to splice variants. The dcDNA-seq dirCLIP approach enabled mapping of SRSF3 binding sites along the full transcript length, often distinguishing even solely 5’ end divergent mRNA isoforms (**Figure 3C-F**). For instance, in the *Nasp* gene, all the crosslink sites mapped to a transcript with an included alternative exon (**Figure 3D**). *Ddx5* encodes an RNA helicase with key functions in RNA metabolism and expresses three isoforms differing in their 5’ UTRs. dirCLIP assigned SRSF3 binding sites exclusively within the shortest 5’ isoform suggesting isoform-specific regulation (**Figure 3E**). Short-read iCLIP mapped SRSF3 binding sites within the last intron of *Tpt1* gene that also carries a small nucleolar RNA gene. With dcDNA-seq dirCLIP, we were confidently able to differentiate between SRSF3 binding to *Tpt1* and *Snora31* transcripts (**Figure 3F**).

**Figure 3.**
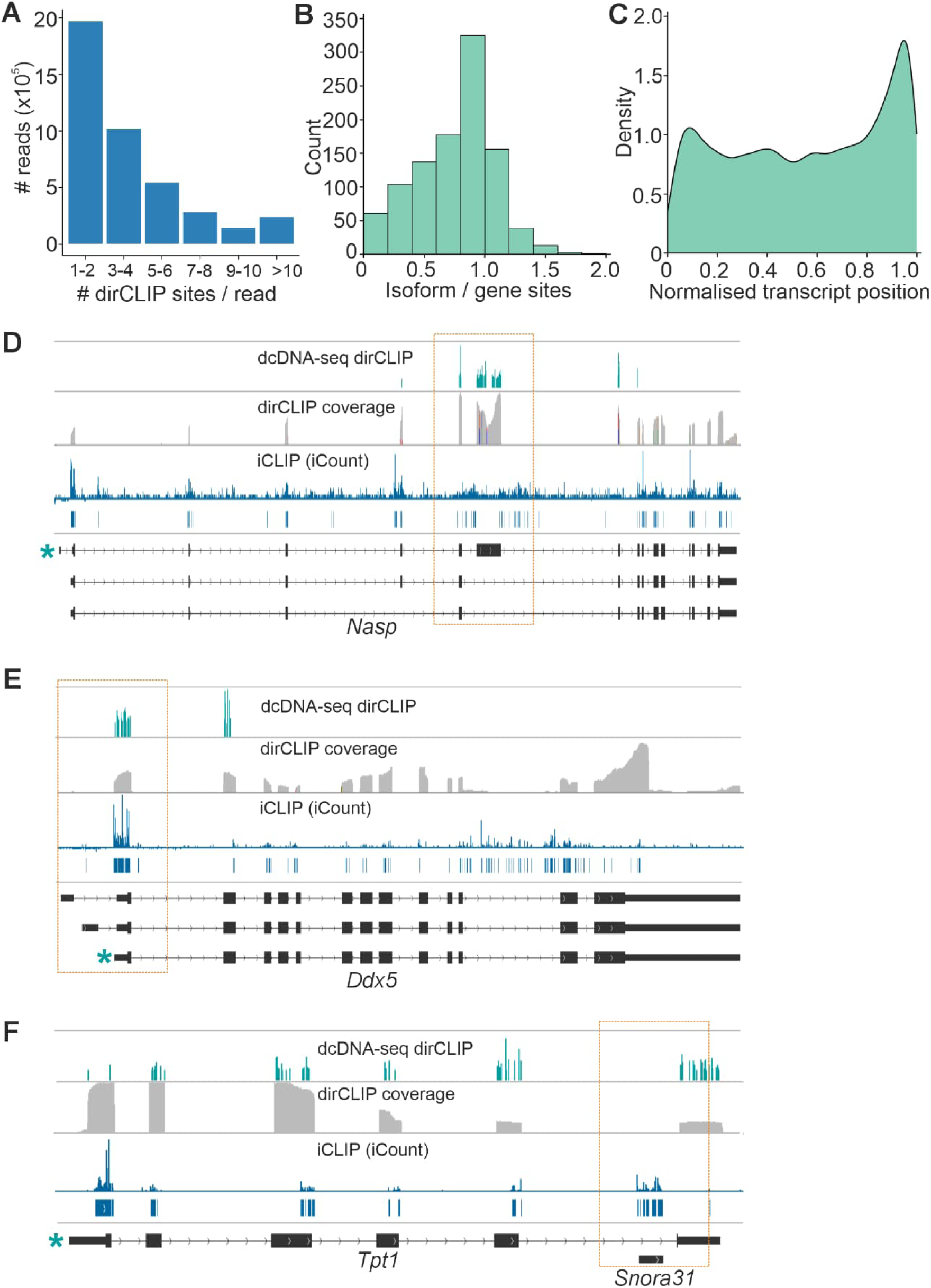
SRSF3 dirCLIP sites map largely to single isoforms. (**A**) Distribution of the number of co-occurring dirCLIP sites per read in one pseudo-replicate. (**B**) Comparison of dcDNA-seq dirCLIP sites determined by genomic and isoform-aware (GENCODE) mutation profiling. The histogram shows the ratio of isoform-level to gene-level sites, with most transcripts showing binding restricted to single isoforms (ratio approaching 1). (**C**) Density distribution of mutation sites across normalized transcript length. (**D-F**) Integrated Genomics Viewer (IGV) tracks of dcDNA-seq dirCLIP sites mapping to the *Nasp*, *Ddx5* and *Tpt1* loci. The upper track shows isoform-aware mapping of dirCLIP sites, and iCLIP peaks are included for comparison. Gene models for the isoforms are shown below. dirCLIP sites were defined using nucleotide coverage ≥50 and a mutation rate difference ≥0.03 and including deletions.

### Discovering the RNA binding prevalence and specificity of HMGA1 with dirCLIP

Having benchmarked dirCLIP with the well-characterized RBP SRSF3 that binds to RNA *via* an RNA recognition motif (RRM), we next evaluated its applicability for discovering new RNA binding activities. High-mobility group protein A1 (HMGA1) is an architectural transcription factor that has been suggested to bind to RNA in human cells ^24–26^. The mouse *Hmga1* gene gives rise to two isoforms, HMGA1a and b, that are intrinsically disordered proteins with three DNA-binding AT-hook motifs and an acidic C-terminal domain (**Figure 4A**). We first determined HMGA1a/b RNA binding capacity in P19 cells by targeted RNA interactome capture that demonstrated binding of both isoforms to poly(A) RNA (**Supplementary Figure 3A**). HMGA1 was also detected in the cellular RNP fraction (**Supplementary Figure 3B**). RNA electrophoretic mobility shift assay (REMSA) using purified HMGA1a/b protein and *in vitro* transcribed stem loop 2 (SL2) of 7SK noncoding RNA (*Rn7sk*), a previously characterized HMGA1 RNA target in human cells, showed direct RNA binding *in vitro* supporting the *in vivo* data (**Figure 4B and Supplementary Figure 3C**) ^24^. Point mutations in the AT-hooks abolished HMGA1 RNA binding both in cells and *in vitro* while deletion of the C-terminal domain had little effect, identifying the AT-hooks responsible for the interaction with RNA (**Figure 4B, Supplementary Figure 3A**). The HMGA1 direct RNA binding *via* the AT-hooks was further confirmed by nucleic magnetic resonance (NMR) (**Figure 4C-F, Supplementary Figure 3D-E**). We observed chemical shift changes in the ^1^H-^15^N HSQC spectra of the amide backbone surrounding the AT-hooks and E-rich C-terminal region upon the addition of 7SK SL2 or an AAUU-rich dsRNA (**Figure 4C & E, Supplementary Figure 3D-E**). Chemical shift changes could occur due to direct nucleic acid binding or intramolecular conformational changes, intrinsically disordered proteins typically showing reduced dynamics upon binding to interaction partners. Heteronuclear ^15^N{^1^H}-NOE of HMGA1a:7SK SL2 complex showed clearly reduced dynamics in the AT-hook regions (positive values), whereas the E-rich C-terminal region remained dynamic (negative values) (**Figure 4D & F**). Together, these results demonstrate that HMGA1 binds to RNA *via* the AT-hooks. Chemical shift changes in the E-rich C-terminal region are in accordance with the release of the dynamic intramolecular interactions with AT-hooks II and III when binding to nucleic acids ^35^.

**Figure 4.**
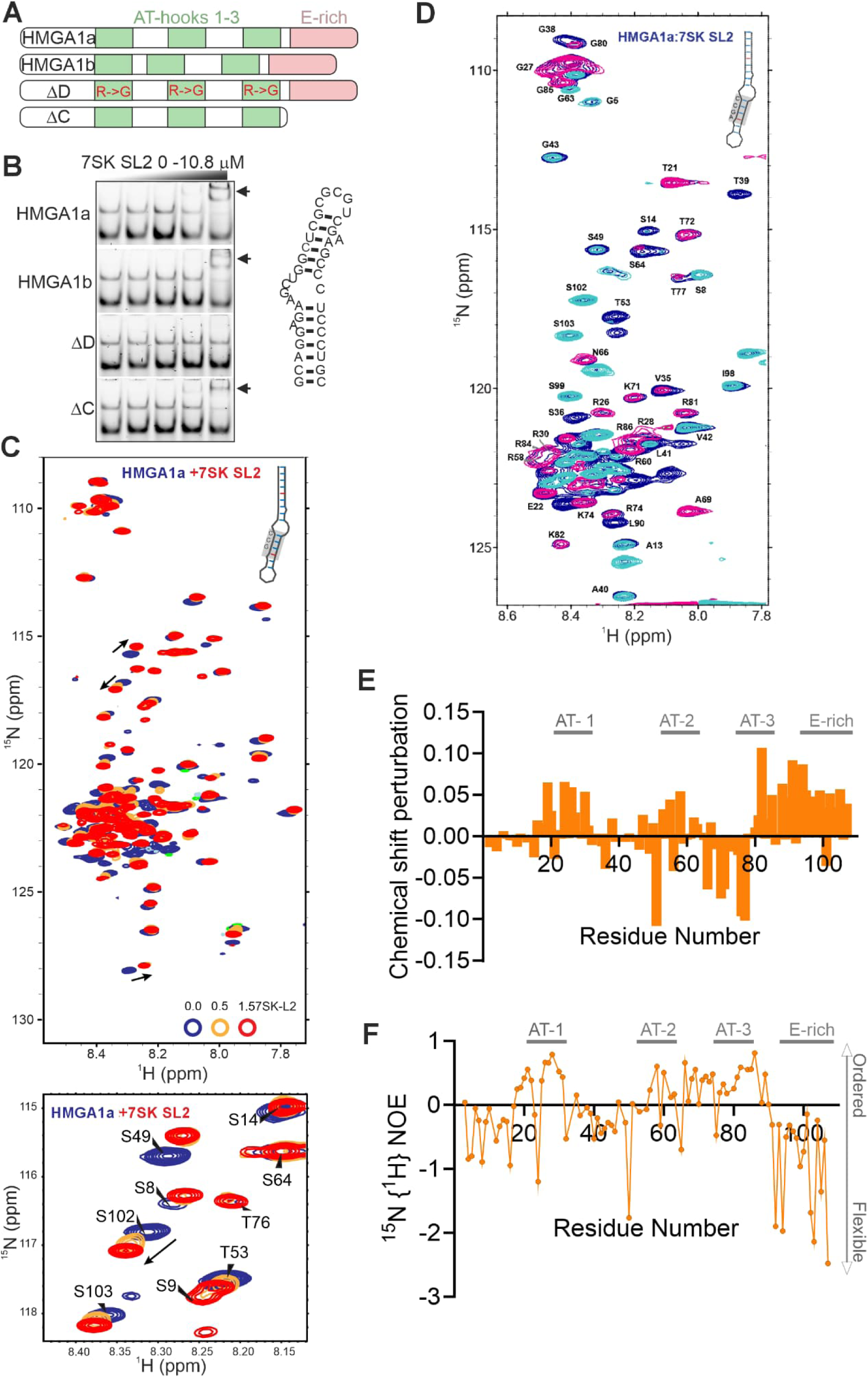
Sequential HMGA1 AT-hook interactions with RNA. (**A**) Schematic representation of the two mouse HMGA1 isoforms a and b (107 and 96 amino acids, respectively), and HMGA1a DNA binding (ΔD) and C-terminal deletion (ΔC) mutants. (**B**) RNA electrophoretic mobility shift assay with mouse 7SK snRNA stem-loop 2 (7SK SL2) and purified HMGA1 proteins. The arrows indicate band shifts. (**C**) ^1^H-^15^N HSQC NMR spectra of free HMGA1a (blue) and HMGA1a bound to 7SK SL2 (yellow, 0.5 molar equivalent; red, 1.5 molar equivalent). The black arrows indicate the chemical shift changes observed when RNA was added, demonstrating RNA interaction. Enlarged sections of the spectra below display chemical shift changes in residues S102 and S49. (**D**) Heteronuclear ^15^H{^1^H}-NOE of HMGA1a:7SK SL2. Reference (blue), positive signals indicating more ordered residues (pink), negative signal indicating dynamic residues (turquoise) (**E**) Combined chemical shift perturbation plots of N-H signals obtained from the ^1^H-^15^N HSQC NMR spectra. (**F**) Intensity ratio from ^15^H{^1^H}-NOE (I_expected_/I_reference_) showing less dynamic residue in the AT-hook regions.

Having demonstrated the RNA binding propensity of mouse HMGA1, we expressed HA-tagged HMGA1a in P19 cells and captured HMGA1a-bound RNAs using an antibody recognizing HA-tag coupled to protein G Dynabeads (**Supplementary Figure 4A-B**). Although dRNA-seq analysis of *in vitro* polyadenylated HMGA1a-bound RNAs detected only few HMGA1a-RNA interaction sites (**Figure 5A, Supplementary Table 3**), we captured the validated RNA target 7SK (**Figure 4**), suggesting that the sites were specific. One of the new putative HMGA1a RNA targets was U1b6 (*Rnu1b6*), an RNA component of the U1 snRNP (**Figure 5A**). HMGA1 has been suggested to interact with U1 small nuclear ribonucleoprotein (snRNP) complex *via* the U1-70K protein component in human cells ^26,36,37^. Unlike in human cells where HMGA1a directly interacts with U1-70K, in mouse P19 cells HMGA1a co-immunoprecipitated with U1-70K only after RNase treatment, suggesting that the protein-protein interaction can occur when RNA bridges are removed (**Figure 5B**). Closer analysis detected putative HMGA1 binding sites in multiple U1 snRNAs at two distinct positions within SL1 and 3 (**Figure 5C-E, Supplementary Figure 4C-E**). Remarkably, dirCLIP detected differential HMGA1 RNA interactions across the four U1 snRNA variants expressed in P19 cells (U1a1, U1b1, U1b2, and U1b6), which differ by only 1-2 nucleotides in length and share >95% sequence identity. While all four variants showed significant HMGA1 binding at position 24 within stem-loop 1 (SL1), variant-specific patterns emerged elsewhere (**Figure 5F, Supplementary Figure 4F, Supplementary Table 4**). REMSA using purified HMGA1a and *in vitro* transcribed SL1 and 3 of U1 snRNA confirmed the AT-hook mediated interaction of HMGA1a with U1 snRNA (**Figure 5G**). Thus, in mouse P19 cells, HMGA1a has mutually exclusive interactions with the RNA and protein components of the U1 snRNP. Our findings demonstrate that dirCLIP can resolve binding differences among highly similar transcript variants that would be indistinguishable by short-read methods, providing unprecedented resolution of RNA-protein interaction specificity. We further observed that HMGA1b interactions with U1-70K and U1 snRNA were weak, suggesting that the two HMGA1 isoforms have differences in their RNA binding activity (**Figure 5B & G**).

**Figure 5.**
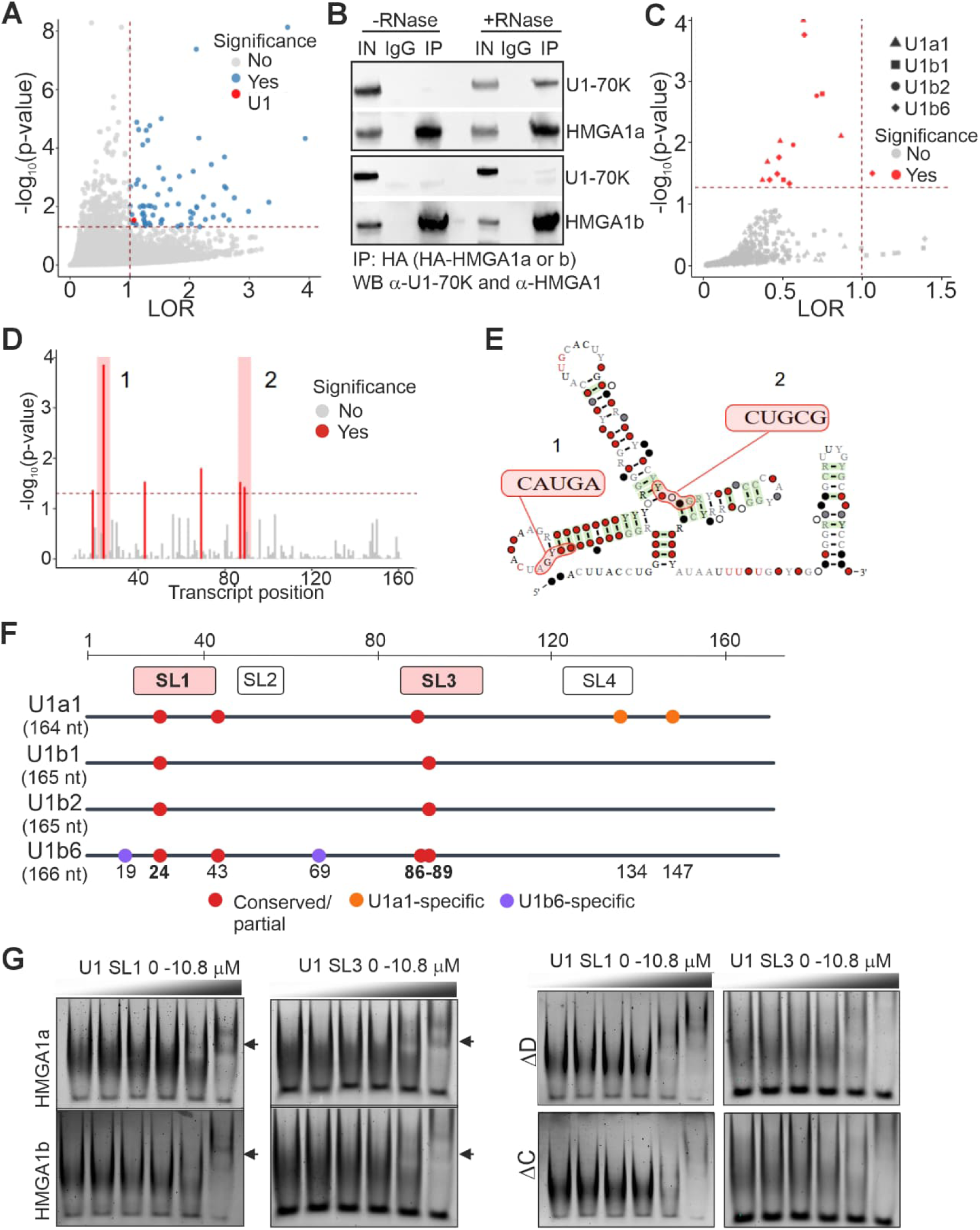
Direct RNA-seq dirCLIP detects HMGA1 binding to U1 snRNA. (**A**) Significant differences in signal intensity and dwell time detected by Nanocompore in dRNA-seq data comparing HMGA1 dirCLIP to untreated transcriptome. (**B**) Co-immunoprecipitation of U1-70K with HMGA1a and b following RNase A-treatment. IN=input, IgG=control, IP=immunoprecipitation. (**C**) dRNA-seq dirCLIP sites meeting significance thresholds in U1 snRNA transcript variants. (**D**) Distribution of dRNA-seq dirCLIP sites meeting significance threshold in U1b6 snRNA variant. Sites marked in pink were detected in multiple U1 snRNA variants. (**E**) Location of the recurring significant dRNA-seq dirCLIP sites in the conserved U1 RNA structure (Rfam ID RF00003, https://rfam.org/family/RF00003). (**F**) Schematic representation of four U1 snRNA variants showing significant HMGA1a binding sites. Structural domains are indicated, SL=stem loop. Circle colors indicate conservation: red, conserved or partially conserved; orange, U1a1-specific; purple, U1b6-specific. (**G**) RNA electrophoretic mobility shift assay with mouse U1 snRNA stem-loop (SL) 1 and 3 and purified HMGA1 proteins. The arrows indicate band shifts. ΔD=HMGA1a with point mutations in the three AT-hooks disrupting DNA/RNA binding; ΔC=C-terminal deletion mutant of HMGA1a.

To enhance the detection of HMGA1 binding sites, we next applied our dcDNA-seq dirCLIP approach to HMGA1a and b. Similar to SRSF3, UV-treated transcriptome (+UV) as the control yielded a smaller number of putative crosslink sites that were a subset of sites detected when applying the untreated transcriptome (-UV) control (**Figure 6A, Supplementary Figure 5A-C, Supplementary Table 5**). We detected substantially more HMGA1a RNA binding sites compared to HMGA1b, and HMGA1b sites largely overlapped with HMGA1a sites. This, together with our *in vitro* data, suggests that the 11 amino acid deletion in HMGA1b may weaken HMGA1 interactions with RNA **(Figure 5G)**. In line with HMGA1 binding preference to AT-rich DNA ^38^ and demonstrating specificity, HMGA1a/b RNA binding sites were AU-rich (**Figure 6B**). Analyzing the global positional patterns of the detected sites, HMGA1a/b sites were found in introns, intergenic regions and coding regions (**Figure 6C**). Mutation profiling by uniquely assigning the reads to transcript isoforms indicated binding to both individual and multiple isoforms with a slight positional preference to the 5’ end (**Figure 6D-E**)

**Figure 6.**
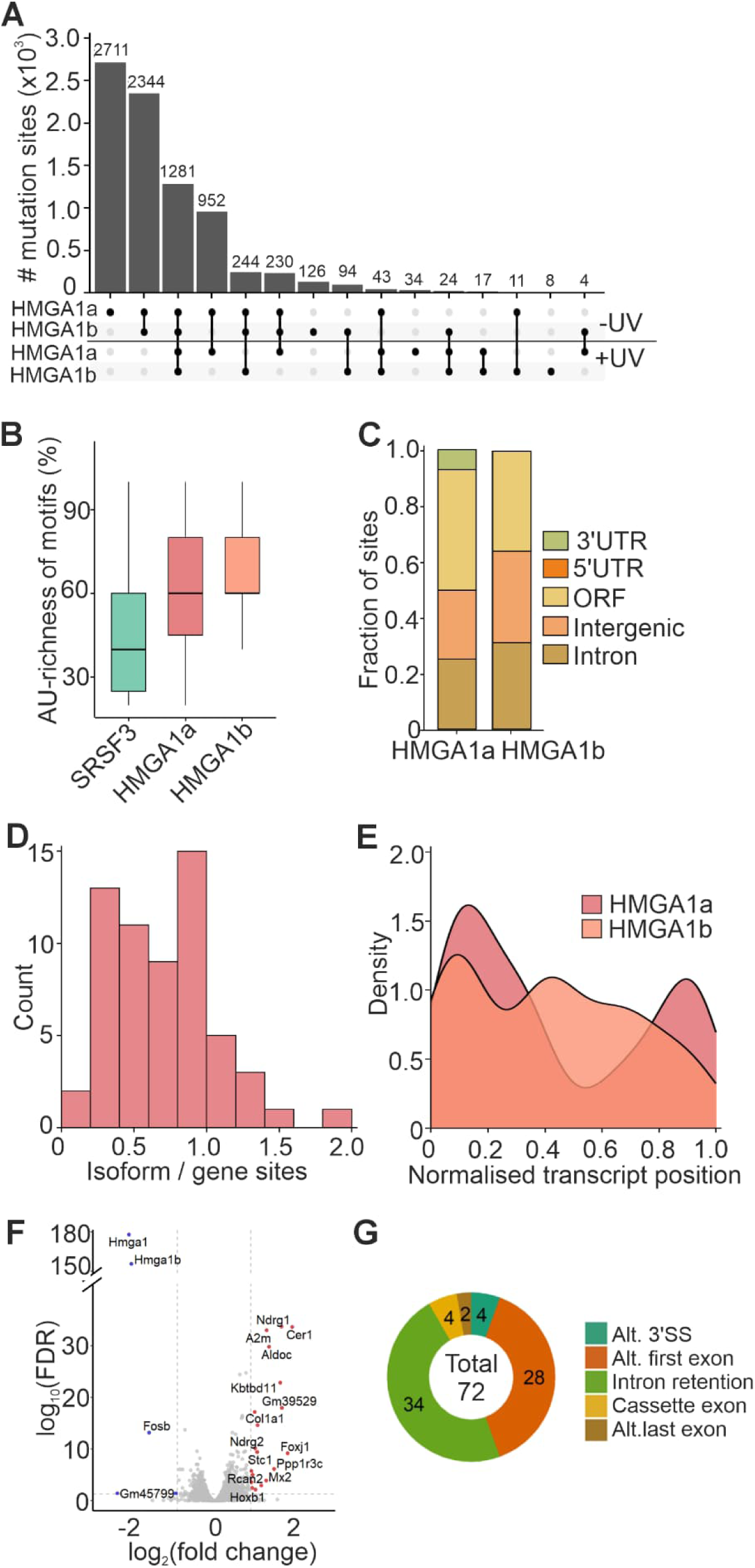
Direct cDNA-seq dirCLIP uncovers HMGA1 RNA binding preference. (**A**) Significant dcDNA-seq dirCLIP sites in HMGA1a and b dirCLIP samples identified using untreated transcriptome (-UV) or UV-treated transcriptome (+UV) controls. Significant dirCLIP sites were defined using a nucleotide coverage ≥50 and a mutation rate difference ≥0.03 including deletions. **(B)** AU-content of the top 50 pentamer motifs based on Z-scores identified within ±50nt of significant SRSF3, HMGA1a and HMGA1b dcDNA-seq dirCLIP sites. Significant dirCLIP sites were defined as in (A). (**C**) Distribution of significant HMGA1a and b dcDNA-seq dirCLIP sites across gene regions. Significant sites were defined as in (A). (**D**) Comparison of dcDNA-seq dirCLIP sites determined by genomic and isoform-aware (GENCODE) mutation profiling. The histogram shows the ratio of isoform-level to gene-level sites. (**E**) Density distribution of mutation sites across normalized transcript length. (**F**) Differentially expressed genes following HMGA1 RNAi in P19 cells. (**G**) Splicing changes following HMGA1 depletion in P19 cells.

To investigate if HMGA1 affected the expression or processing of its target RNAs, we knocked down *Hmga1* using RNAi and conducted short-read RNA sequencing (**Supplementary Figure 5D**). Surprisingly, HMGA1 downregulation led to only minor gene expression and alternative splicing changes in P19 cells (**Figure 6F-G, Supplementary Table 6 and 7**). These findings were supported by ENCODE HMGA1 knockdown data in human cells (**Supplementary Figure 5E-F, Supplementary Table 8 and 9**). Since HMGA1a interacts with U1 snRNP and thousands of other RNAs without affecting their abundance or splicing, HMGA1-RNA interactions may reflect riboregulation, an emerging concept in cell biology ^39,40^.

## DISCUSSION

We have developed dirCLIP, a method that leverages nanopore direct long-read sequencing for mapping of RNA-protein interactions in full-length transcripts. By preserving native transcript context without amplification or fragmentation, dirCLIP enables the mapping of RBP sites in distinct transcript variants. Compared to established methods such as iCLIP and eCLIP, dirCLIP offers unique advantages including full-length transcript coverage, detection of co-occurring binding events in single RNAs and isoform-level resolution, albeit with current throughput limitations (**Supplementary Table 10**). Application of dirCLIP to SRSF3 revealed predominant binding to specific mRNA isoforms that has remained invisible to conventional short-read CLIP methods. The ability to distinguish binding patterns among near-identical transcript variants was further exemplified by HMGA1 interactions with U1 snRNA. The variant-level resolution of RBP mapping has profound implications for understanding RNA regulation as it suggests RBPs may differentially regulate distinct splice variants and closely related transcript variants to achieve regulatory complexity.

The two detection strategies employed in dirCLIP, dRNA-seq coupled with signal perturbation analysis and dcDNA-seq with mutation profiling, captured overlapping sets of crosslinked RNA-protein complexes. In dRNA-seq detection, signal deviation was complemented with read termination analyses to detect different, largely non-overlapping crosslink signatures. Signal changes likely reflect smaller adducts or partial crosslinking events that permit RNA translocation through the nanopore, while termination events indicate more substantial modifications that physically obstruct pore passage. This complementarity provides orthogonal validation of binding sites and offers insights into the heterogeneous nature of UV-induced crosslinks, which may vary depending on the local RNA structure, protein orientation, and crosslinking efficiency at different sites, features that can be further explored ^27,41–43^. Recent advances in detecting endogenous RNA modifications through shifts in current signal and dwell times validate our approach of using biophysical signatures to detect protein crosslinks ^44–46^.

The pronounced 3’ bias observed in dRNA-seq reflects both the inherent directional nature of nanopore sequencing and the compound effect of crosslink-induced termination. While the Nanocompore algorithm ^28^ we employed has proven effective for detecting RNA modifications by comparing modified and unmodified samples, its application to protein crosslinks revealed unique challenges. The larger size and heterogeneous nature of protein adducts compared to chemical modifications likely contribute to the distinct signature patterns we observed. Development of larger pores enabling more efficient translocation of nucleotides with bigger adducts and/or future improvements in basecalling algorithms specifically trained on crosslinked RNA, similar to recent developments for detecting N6-methyladenosine (m6A) and other modifications ^47,48^, could enhance detection sensitivity and expand coverage toward 5’ regions.

The two detection strategies employed in dirCLIP, dRNA-seq coupled with signal perturbation analysis and dcDNA-seq with mutation profiling, captured overlapping sets of crosslinked RNA-protein complexes. The superior performance of dcDNA-seq over dRNA-seq for comprehensive binding site detection reflects fundamental differences in how these approaches handle crosslinked RNA. The ability of reverse transcriptases, particularly MMLV-based enzymes, to read through protein adducts while introducing “diagnostic” mutations has been successfully exploited for RNA structure probing ^29,31,32^. Our finding that deletions constitute a significant proportion of crosslink-induced mutations and improve detection specificity when included in the analysis challenges the conventional view that deletions frequently represent sequencing errors. This observation aligns with structural studies showing that reverse transcriptases can skip damaged or modified nucleotides ^49^, suggesting deletion signatures may be particularly informative for identifying bulky amino acid adducts.

Our benchmarking with SRSF3 demonstrated that dirCLIP achieves comparable specificity to the established iCLIP method while providing unprecedented isoform resolution. The >75% concordance with validated CNNC motifs within fifty nucleotides of the detected sites confirms that dirCLIP captures genuine binding events rather than artifacts. This was further supported by the overlapping sites detected with the dRNA- and dcDNA-seq dirCLIP modalities. The finding that SRSF3 binding sites predominantly localize to single isoforms rather than being shared across splice variants provides key new insights into SR protein function. The isoform specificity may explain how SRSF3 can promote inclusion of some exons while simultaneously facilitating skipping of others within the same transcript ^50^, a paradox that has puzzled the splicing field.

The application of dirCLIP to HMGA1 demonstrates the method’s utility for characterizing noncanonical RBPs lacking well-defined RNA recognition domains ^39,40^. Our biochemical validation confirmed that the DNA binding AT-hooks ^38^ also mediate RNA interactions. The NMR analysis revealing dynamic interactions of the three AT-hooks with RNA provides mechanistic insight into how intrinsically disordered proteins can achieve RNA recognition despite lacking structured RNA binding domains. HMGA1 dirCLIP demonstrated the ability to detect specific binding to distinct RNA variants and revealed new AT-hook dependent RNA interactions. The mouse genome encodes multiple U1 variants (U1a1, U1b1, U1b2, U1b6) that differ by only 1-2 nucleotides yet showed distinct HMGA1 binding signatures. While a conserved binding site in stem-loop 1 was detected across all variants, several positions showed variant-specific binding, including sites unique to U1a1 or U1b6. Such discrimination would be impossible with fragmented short-read approaches, where reads from different U1 variants would be ambiguously mapped or collapsed together. The observation that HMGA1 depletion causes minimal changes in gene expression or splicing, despite widespread RNA binding, suggests HMGA1-RNA interactions may serve regulatory functions beyond conventional RBP activities. This finding aligns with emerging concepts of riboregulation, where RNAs act as allosteric regulators or scaffolds that modulate protein activity or localization ^39,40,51^.

The current limitations of dirCLIP primarily relate to sequencing throughput constraints of existing nanopore platforms. While the transition to PromethION capable of generating >100 Gb per flow cell substantially increases coverage depth ^52^, achieving the coverage depth typical of short-read CLIP experiments remains challenging. However, Oxford Nanopore’s rapid technology development trajectory, with consistent improvements in output and chemistry, suggests this gap will continue to narrow. Importantly, the unique biological insights gained from preserving full-length transcripts - particularly RNA variant-specific binding patterns - justify the current throughput trade-off for many applications. The recent chemistry update with improved basecalling accuracy should further enhance both dRNA- and dcDNA-seq detection sensitivity.

Several technical improvements could expand dirCLIP capabilities. Integration of RNA modification detection algorithms ^45,53,54^ could enable simultaneous mapping of RBP binding sites and epitranscriptomic marks, revealing potential crosstalk between these regulatory layers. Transfer learning approaches recently developed for detecting multiple RNA modification types ^55^ could be adapted for crosslink detection, potentially reducing the amount of training data required for new RBPs. Incorporation of barcoding strategies would enable multiplexed analysis of multiple RBPs or conditions within a single sequencing run, improving cost-effectiveness and enabling direct comparative analyses ^56–58^.

dirCLIP opens several exciting research avenues. Application to single-cell or low-input samples, enabled by continued improvements in nanopore sensitivity, could reveal cell-type-specific RBP-RNA interaction networks. Integration with spatial transcriptomics approaches could map RBP binding patterns within tissue contexts. The ability to detect binding sites across full-length transcripts and within single RNA molecules also enables investigation of long-range cooperative or competitive interactions between multiple RBPs on the same RNA molecule, providing insights into the combinatorial RBP code that governs post-transcriptional gene regulation ^17,59,60^. The ability to map isoform-specific RBP sites has important implications for understanding disease mechanisms and developing therapeutics. Many disease-associated mutations affect splicing patterns ^61–63^, potentially disrupting isoform-specific RBP interactions that conventional CLIP methods cannot detect. For therapeutic development, our results suggest that targeting specific RNA isoforms rather than all transcripts from a gene may achieve more precise modulation of RBP-RNA interactions. This is particularly relevant for antisense oligonucleotide and small molecule approaches targeting RNA-protein interactions where isoform selectivity could reduce off-target effects. In neurodegenerative diseases where toxic protein isoforms drive pathology, dirCLIP could identify isoform-specific RBP binding sites that serve as precision therapeutic targets, preserving beneficial isoforms while selectively modulating pathogenic variants. The ability to detect binding patterns in full-length transcripts enables rational design of splice-switching therapeutics that consider the complete regulatory context, including long-range RBP interactions and combinatorial binding events invisible to fragmentation-based methods.

Several limitations of dirCLIP should be considered when designing experiments. First, nanopore sequencing throughput, while rapidly improving, remains lower than short-read platforms, requiring careful experimental design to achieve sufficient coverage depth for comprehensive binding site detection. Current experiments typically yield 1-5 million mapped reads per flow cell, compared to tens of millions achievable with Illumina-based CLIP methods. This throughput constraint is most pronounced for dRNA-seq, where crosslink-induced termination further reduces coverage of 5’ transcript regions. Second, the sensitivity of crosslink detection depends on the nature of the RBP-RNA interface. RBPs with extensive RNA contacts may generate more detectable crosslink signatures than those with limited interaction surfaces. Similarly, the efficiency of UV crosslinking varies with RNA structure and sequence context, potentially biasing detection toward certain binding sites. Third, while dcDNA-seq provides more uniform transcript coverage than dRNA-seq, it requires reverse transcriptase read-through of protein adducts, which may not occur efficiently for all crosslink types. The balance between read-through (enabling mutation profiling) and termination (limiting upstream detection) likely varies among RBPs and crosslinking conditions. Fourth, isoform-level analysis requires sufficient read depth per transcript variant, limiting detection to moderately or highly expressed isoforms. Lowly expressed splice variants may not achieve the coverage thresholds required for confident binding site calls. Finally, computational analysis of long-read data remains more resource-intensive than short-read pipelines, and standardized tools for RBP site detection from nanopore data are still emerging. We anticipate that continued development of basecalling algorithms, signal analysis methods, and reference-guided approaches will address many of these limitations.

In conclusion, dirCLIP represents a fundamental advance in our ability to interrogate RNA-protein interactions by preserving the native transcript context essential for understanding isoform-specific regulation. As nanopore sequencing technology continues to mature, with improvements in throughput, accuracy, and modification detection capabilities, dirCLIP and related approaches will become increasingly powerful tools for deciphering the complexity of post-transcriptional gene regulation.

## DATA AND CODE AVAILABILITY

Sequencing data has been submitted to Sequence Read Archive (SRA) under BioProject ID: PRJNA1304717.

Custom code has been submitted to GitHub (https://github.com/rnaomics/dirCLIP).

## ACKNOWLEDGEMENTS

We thank Eduardo Eyras and Aditya Sethi for helpful discussions in the early phases of this project and are grateful to Alice Cleynen for insights and guidance with the statistical analyses. The authors are grateful to the representatives of Oxford Nanopore Technologies for insightful discussions and technical advice. This work was supported by the Victorian Government Operational Infrastructure Support Scheme [to MLÄ], National Health and Medical Research Council (NHMRC) [GNT1138870 to MLÄ and JK], Sigrid Juselius Foundation [to MLÄ], Jane and Aatos Erkko Foundation [to MLÄ], Finnish Cultural Foundation/Pirkanmaa Fund [to MS], Research Council of Finland [to MLÄ], NHMRC Investigator Grant [GNT1175388 to NES], Bootes Foundation Grant 2022 [to NES], Australian Research Council (ARC) Discovery Grants [DP180100111 and DP250103133 to NES], and CNRS Postes Rouges 2025 grant [to NES].

## AUTHOR CONTRIBUTIONS

MS, ZZ, MR and FL conducted the experimental work. ZZ, MS and KW prepared sequencing libraries and conducted sequencing. KW conducted computational analyses related to direct RNA sequencing, MB to direct cDNA and TA to short-read sequencing. MLÄ and NES conceptually designed the project and MLÄ, NES and JK supervised the work. MLÄ compiled the manuscript with significant contributions from NES, KW, ZZ, TA, MS and MB. All authors read and approved the manuscript.

## METHODS

### Cell lines and constructs

Murine P19 embryonic teratocarcinoma and HEK293 cells were maintained in High-glucose Dulbecco’s Modified Eagle Medium supplemented with 2mM GlutaMAX, 50U/ml penicillin/streptomycin and 10 % fetal bovine serum. The cells were grown on 0.1% gelatine-coated dishes at 37°C with 5% CO_2_. The P19 cells expressing EGFP-tagged SRSF3 from a BAC-transgene (SRSF3-BAC) were a kind gift from Michaela Müller-McNicoll ^22^.

Four vectors were constructed for transient expression of HMGA1 protein variants with an EGFP-reporter. The vectors contained the sequences encoding HA-tagged versions of mouse HMGA1 isoforms a or b, or HMGA1a DNA binding (DD) or C-terminal deletion (DC) mutants fused with a T2A self-cleaving peptide and EGFP. HMGA1a ΔD carried a point mutation substituting the second arginine to glycine in each of the three P-R-G-R-P conserved AT-hook motifs (positions 28, 60 and 86), and the HMGA1a ΔC had a 19 amino acid (57 base pair) deletion at the C-terminus. P19 wildtype or HEK293 cells were transfected with the vectors using TurboFect™ Transfection Reagent (Thermo Fisher Scientific) according to the manufactureŕs recommendations.

For bacterial expression, the codon optimized sequences encoding the HMGA1 variants were cloned into pETGB_1a bacterial expression vectors. An N-terminal hexa-histidine purification tag (HIS6) and Streptococcal protein G (GB1) were added to increase solubility and yield. A TEV protease cleavage site was included to enable cleavage of the HIS6-GB1.

### UV crosslinking and immunoprecipitation

P19 SRSF3-BAC cells or P19 wildtype cells transiently expressing HA-tagged HMGA1 grown on monolayer were crosslinked on ice at 200 mJ/cm^2^ and harvested. The cell pellet was resuspended in CLIP lysis buffer (50mM Tris-Cl pH 7.4, 100mM NaCl, 1% Igepal, 0.1% SDS and 0.5% sodium deoxycholate) supplemented with protease inhibitors (Merck) and 1:1000 RNase inhibitor (40U/µl, Thermo Fisher Scientific) and incubated for 15min on ice. The lysate was mildly sonicated and treated with TURBO DNase I (Thermo Fisher Scientific) to facilitate cell lysis and chromatin solubility. The cleared lysate was incubated at 4°C with GFP-Trap magnetic agarose beads (Chromotek) for SRSF3-EGFP and with Dynabeads-Protein G (Thermo Fisher Scientific) coupled with anti HA-tag antibody (Invitrogen, HA-C5) for HA-HMGA1a/b. The beads were washed thrice with high-salt wash buffer (50mM Tris-Cl pH 7.4, 1M NaCl, 1% Igepal, 0.1% SDS and 0.5% sodium deoxycholate) and twice with PNK buffer (20mM Tris-Cl pH 7.4, 10mM MgCl_2_, 0.2% Tween-20). RNA-bound proteins were digested with Proteinase K (Merck) in PK buffer (100mM Tris-Cl pH 7.4, 50mM NaCl and 10mM EDTA) for 20min at 37°C 1100rpm, followed by incubation in PK buffer supplemented with 7M urea at 50°C for 20min 1100rpm. RNA was extracted from the supernatant with phenol:chloroform:isoamyl alcohol (125:24:1, pH 4.5). Similarly, RNA was isolated from untreated and UV-treated P19 SRSF3-BAC and wildtype cells. The dirCLIP and total RNA samples were used for sequencing library preparation using the native poly(A) tail or *in vitro* polyadelylated using an *E. coli* Poly(A) Polymerase (New England Biolabs) with a modified protocol to add ∼100nt of poly(A). The total RNA samples were ribodepleted before polyadenylation (NEBNext rRNA Depletion kit, New England Biolabs).

### Library preparation and nanopore direct long-read sequencing

The library preparation for direct RNA sequencing was performed using an SQK-RNA002 kit (Oxford Nanopore) according to the manufacturer’s instructions and as previously described ^45,64^. The RNA control standard was excluded from the reactions. Following reverse transcription and motor protein ligation, libraries were purified using Agencourt RNAClean XP beads (Beckman Coulter). Sequencing was performed using R9.4.1 flow cells and the MinION Mk1B device according to the manufacturer’s instructions. Flow cells were equilibrated to ambient temperature (25°C) for 30min before use and verified to contain >800 active pores. Data acquisition was controlled through MinKNOW software (version 4.3.25) with bulk FAST5 output enabled for downstream signal analysis. Sequencing runs proceeded for 24-72h depending on pore availability and throughput. Real-time base calling was performed using the integrated Guppy basecaller.

Direct cDNA sequencing libraries were prepared using the reverse transcription and strand-switching method utilizing Maxima H Minus reverse transcriptase and the Ligation Sequencing Kit V14 (Oxford Nanopore Technologies SQK-LSK114). For the manganese buffer conditions, magnesium chloride in the buffer disclosed by the manufacturer was replaced with the same amount of manganese chloride ^33^. The cDNA was sequenced on MinION Flow Cells (R10.4.1) using a MinION Mk1B sequencing device as described above. Sequencing runs proceeded for 24-72h depending on pore availability and throughput. Real-time base calling was performed using the integrated Guppy basecaller.

### Data processing of dRNA-seq dirCLIP

Fast5 files generated from sequencing were basecalled using Guppy 5 with the parameters --flowcell FLO-MIN106 and --kit SQK-RNA002. Basecalled fastq files were then mapped to the ENSEMBL *Mus Musculus* GRCm39 reference transcriptome. SRSF3 datasets were mapped to the cDNA reference fasta, whilst the HMGA1a datasets were mapped to the combined cDNA and ncRNA reference fasta to capture non-natively polyadenylated RNAs such as snRNAs. Mapping was conducted using minimap2 with the parameters -ax map-ont - k14 -uf. samtools view -F 0×904 was used to filter the mapped files for primary alignments, followed by sorting and indexing.

Nanopolish index was used with sequencing summary, fast5 and fastq file inputs to generate an index file of read IDs and their corresponding fast5 file paths. Nanopolish eventalign was then used with the options --scale-events --print-read-names --samples to align signal information with the reference fasta files used in the mapping above, generating an output of signal events across transcript positions. The event level information was then collapsed as a summary at each transcript position using NanopolishComp Eventalign_collapse. Conditions were then compared using Nanocompore sampcomp with the parameters --mincoverage 5, --comparison_methods GMM and --max_invalid_kmers_freq 0.6 specified. Significant sites were then derived using a |LOR| threshold >1 and an adjusted p-value <0.005 for the SRSF3 datasets, with a lower significance threshold of 0.05 used for the HMGA1a datasets due to reduced coverage. For datasets that had greater than 500,000 reads, a single replicate was randomly split into three pseudo-replicates after Nanopolish, using the read ID information stored in the Nanopolish output.

Bedtools bamtobed was used to extract read IDs, read start and read end coordinates of mapped transcripts. For the datasets split into technical replicates, the extracted read IDs from Nanopolish were used to divide the read coordinate files into the same segmented read groups, ensuring uniformity of replicates between the Nanocompore and read termination analyses. From the output bed files, the number of reads ending at each transcript coordinate were counted and counts matrices were generated including all replicates of test conditions. Prior to significance testing, positions with less than ten reads were removed. A Poisson GLM model was used to determine significance, with significance thresholds of log_2_ fold change >2 and adjusted p-values <0.05 applied.

Sites that occurred within 5nt of each other were collapsed together into a single site region, taking the range center as the apex of the crosslinking event. The highest log_2_ fold change or LOR value and lowest adjusted p-value was included as a summary statistic for that region.

The package Biostrings in R was used to generate fasta files of 50nt ranges around the collapsed significant dRNA-seq sites. Frequency of motifs was compared to a randomized background of the same sequences shuffled hundred times, with significance assessed by z-score and Fischer’s exact tests. Sequence logo plots were generated on motifs with z-score >2 and p-value <0.05 using the R package ggseqLogo.

Salmon quant ^65^ with standard parameters was used to estimate transcript expression levels in the untreated and UV-treated transcriptomes. Lowly expressed transcripts were removed by implementing a cutoff of TPM > 0.5. BiomaRt was used to add gene-level information to each transcript, and the output was further filtered to include only genes that exhibited statistically significant dirCLIP sites in the SRSF3/+UV or SRSF3/-UV comparisons. The number of isoforms per gene was then compared to the number of isoforms with binding sites per gene, with any instances of greater than or equal to 5 transcripts grouped.

The R packages GenomicFeatures and GenomicRanges were used to map transcript coordinates of the dRNA-seq sites to the genome using an ENSEMBL GRCm39 gtf file as reference. Significant cDNA-seq sites were converted to mm39 genomic coordinates from mm9 coordinates using the UCSC lift over tool.

### Data processing of dcDNA-seq dirCLIP data

Direct cDNA sequencing reads were base called using Dorado v0.3.4 with the model dna_r10.4.1_e8.2_400bps_sup@v4.2.0. The reads were aligned to the *Mus musculus* mm9 reference genome using Minimap2 v2.24-r1122 ^66^ with the parameters -ax splice -uf -k14. Secondary and supplementary alignments were removed with SAMtools v1.19 ^67^ retaining only primary alignments. Bam files were converted to bed format using BEDTools v2.31.0 ^68^ for downstream analyses. Reads <20nt were excluded from the analysis. For violin plot visualizations, reads exceeding the 99th percentile of the read length distribution were removed. To improve nucleotide coverage, the bam files from the two biological replicates of the SRSF3 dirCLIP samples were merged using samtools merge. The untreated transcriptome sample consisted of a single replicate of sufficient read depth. Pseudo-replicates were generated by randomly subsampling 70% of the reads from each merged bam file using samtools view with three different random seeds. This subsampling approach ensured consistency in read depth across analyses while preserving biological variability.

Bam files were filtered to remove reads with a mean base quality score below 8 as low-quality reads could inflate mutation rates. Per-nucleotide mutation rates were then calculated using Perbase v0.9.0 with a base quality filter (-Q 8). Bases with a quality score below 8 were masked and represented as ‘N’. Mutation rates were calculated both including and excluding deletions. Insertions were not considered in the mutation rate calculation and positions with masked ‘N’ bases were excluded. Only nucleotides that met the specified coverage threshold in all pseudo-replicates of both the dirCLIP and corresponding control samples were retained. Coverage thresholds of 50, 100, 200, 300, 400, and 500 were evaluated.

The mean mutation rate at each nucleotide position was computed from the pseudo-replicates. For dirCLIP samples, background mutation rates were subtracted using either untreated (-UV) or UV-treated (+UV) transcriptomes as controls. A two-sample t-test was performed to evaluate whether mutation rates in dirCLIP samples were significantly higher than in the corresponding control samples. To account for multiple testing, p-values were adjusted using the Benjamini–Hochberg (BH) correction method. Sites with an adjusted p-value <0.05 were considered statistically significant. To further reduce false positives, a mutation rate difference threshold of 0.01, 0.02, 0.03, 0.04, 0.05, or 0.06 were evaluated. A nucleotide was considered a significant mutation site if it met all three criteria:

1. Coverage above the coverage threshold,
2. Adjusted p-value <0.05,
3. Mutation rate difference greater than the mutation rate difference threshold.

Isoform-specific read assignment was performed using Bambu version v3.10.0 ^69^ with the discovery = FALSE. Bam files aligned to the mm9 reference genome and GENCODE GTF were provided as input to Bambu. Reads mapping to multiple isoforms were excluded from downstream analysis. After assigning reads to isoforms, read sequences were aligned to isoform-specific fasta files using Minimap2 ^66,70^ with parameters -ax splice -uf, generating separate bam files for each transcript isoform. Bam files from biological replicates were merged, subsampled to generate pseudo-replicates, per-nucleotide mutation rates were calculated, and significant mutation sites were identified as above. Sites located within ±5nt of each other were merged into single continuous regions to avoid overcounting closely spaced events.

The co-occurrence of dirCLIP sites within individual sequencing reads was determined by first adding MD tags to each pseudo-replicate bam file using samtools calmd to enable identification of base mismatches and deletions relative to the reference sequence. The positions of base mismatches and deletions in the reference sequence were then extracted from individual reads using the pysam Python library. For each read, the positions of all mutation events (mismatches and deletions) were recorded in reference transcript coordinates. These events were then filtered to retain only those overlapping significant dirCLIP sites (nucleotide coverage ≥50 and a mutation rate difference ≥0.03) and the co-occurring dirCLIP sites per read were counted.

### Data processing of iCLIP data and comparison to dirCLIP

Raw iCLIP sequencing data for three SRSF3 iCLIP and four control samples were downloaded from the GEO database (accession number: GSE69689). Replicates for each condition were merged using the cat command. The merged fastq files were processed using the nf-core/clipseq pipeline v1.0.0 (peakcallers iCount, Paraclu, Pureclip, Piranha) and reference genome *Mus musculus* mm9 (mm9.fa) with the annotation gencode.vM1.annotation.gtf ^71^. The iCount peaks were used for downstream analysis.

The overlap between significant SRSF3 dirCLIP mutation sites and SRSF3 iCLIP peaks (identified using the iCount peak caller) was assessed by defining a ±5nt window around each dirCLIP site and testing for intersection with iCLIP peak regions. Enrichment of pentamer motifs was assessed in the vicinity of SRSF3 iCLIP peaks and dirCLIP sites by extracting a 100nt window (±50nt) from the peak/site center in the mm9 reference genome. Within each window, all possible pentamers were enumerated using a sliding window approach, and their frequencies were calculated. Background distributions were generated by randomly shuffling the nucleotide sequences within each window hundred times while preserving local nucleotide composition. Pentamers were extracted from these shuffled sequences using the same sliding window approach. For each pentamer, the mean and standard deviation of its count across the hundred shuffled sequences were calculated. To quantify enrichment, Z-scores were computed for each pentamer. Statistical significance of enrichment was evaluated using Fisher’s exact test. Pentamers were ranked first by Z-score, followed by ranking according to Fisher’s test p-values, to identify the most enriched motifs. For SRSF3 iCLIP and dirCLIP data, the shortest distance between each iCLIP peak center/dirCLIP site and the nearest SRSF3-binding CNNC motif was calculated. As a control, the nucleotide sequences within a 100nt window (±50nt from the peak center) were randomly shuffled three times. For each shuffled sequence, the distance between its center and the nearest CNNC motif was determined.

### Protein production and *in vitro* transcription

pETGB1_HMGA1constructs were transformed into *E. coli* BL21(DE3) Rosetta 2 cells and grown on LB medium supplemented with 50μg/ml kanamycin and 34μg/ml chloramphenicol and protein expression was induced with 1.5mM isopropyl-β-D-1-thiogalactosid (IPTG) and expressed at 20^°^C for 24h. Bacterial pellets were washed with RBS buffer (10mM HEPES pH 7.4, 10mM NaCl, 3mM MgCl_2_) before lysis (50mM KPO_4_, 1M NaCl, 5mM imidazole, 1mM β-mercaptoethanol 1% Triton X-100, lysozyme, DNase I and 5mM MgCl_2_ and Protease Inhibitors (Roche)). The HIS6-tag proteins were purified by binding to Ni-NTA (nickel-nitrilotriacetic acid) beads (Qiagen) at 4°C. Beads were washed by gravity flow twice with 50mM K_2_HPO_4_/KH_2_PO_4_ pH 7.6, 1M NaCl, 10mM imidazole and once with 50mM K_2_HPO_4_/KH_2_PO_4_ pH 7.2, 150mM NaCl, 30mM imidazole. HIS6-tag proteins were eluted with 50mM K_2_HPO_4_/KH_2_PO_4_ pH 7.8, 100mM NaCl, 150mM imidazole, concentrated and passed through two HiTrap Heparin HP affinity columns (GE biosciences) using an Akta Purifier (GE Healthcare Life Sciences) to separate truncated proteins and any bound RNA/DNA. Columns were pre-equilibrated with buffer A (20mM K_2_HPO_4_/KH_2_PO_4_ pH 7.2, 0.5mM EDTA, 1mM DTT), and the proteins eluted with buffer B (20mM K_2_HPO_4_/KH_2_PO_4_ pH 7.2, 1.5M NaCl, 0.5mM EDTA, 1mM DTT) with a gradient of 0.15M to 1M NaCl.

ssDNA oligonucleotides encoding 7SK snRNA SL2, U1 snRNA SL1 or SL3 or an AAUU-rich dsRNA (PRDII) and containing T7 promoter sequence (Integrated DNA Technologies or Sigma-Aldrich) were annealed, and *in vitro* transcription was performed either with T7 polymerase produced in-house or Invitrogen™ MEGAscript™ T7 Transcription Kit (Thermo Fisher Scientific). The samples were treated with 2U TURBO DNase (Thermo Fisher Scientific) and purified using acidic phenol:chloroform, pH 4.5 and isopropanol precipitation. RNA samples were analyzed by running on Agilent Tape Station automated electrophoresis system (Agilent Technologies, Santa Clara, CA, USA).

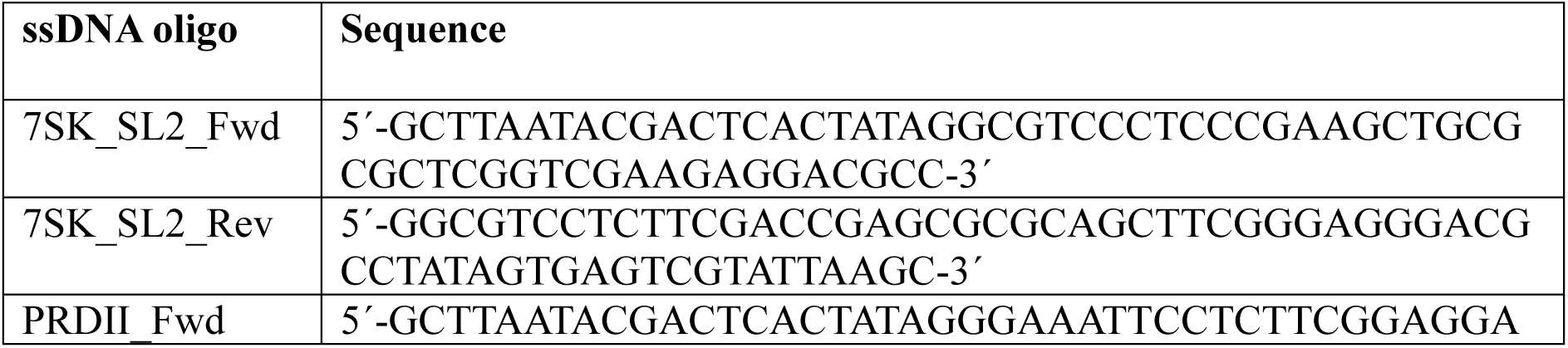
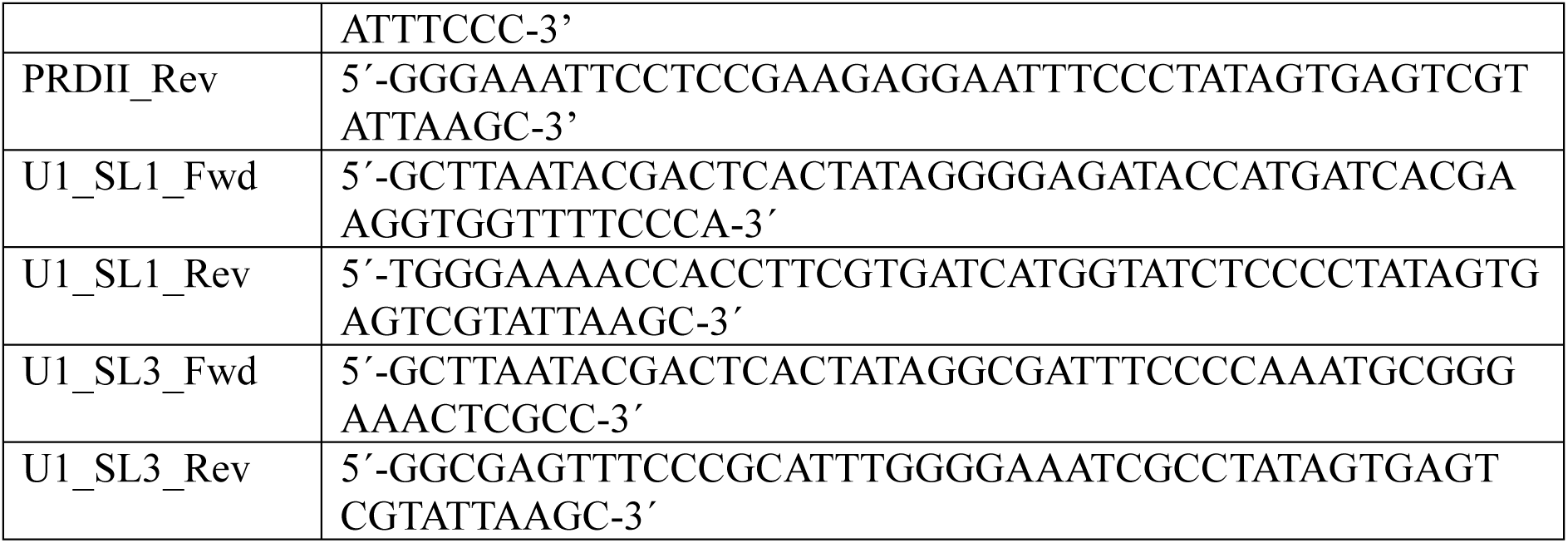

### RNA electrophoretic mobility shift assay (REMSA)

*In vitro* transcribed RNAs were folded by heating at 95°C for 2min and then cooling on ice-water slush for 2min. Increasing concentrations of each protein (0 to 10.8µM) were used to assemble binding reactions with 0.4µM of RNA in Gel shift buffer (20mM HEPES pH 7.2, 100mM KCl, 10mM MgCl_2_, 10% glycerol, 1mM DTT) supplemented with 0.03µg/ml heparin. Each reaction was incubated at 25°C for 15min and subsequently run on a 6% native acrylamide gels in 0.5x TBE. The gels were stained with SYBR Gold (Thermo Fisher Scientific) or GelStar® (Lonza Bioscience) nucleic acid gel before imaging.

### NMR Spectroscopy

To ^15^N and ^13^C isotopic label HMGA1a protein, *E. coli* BL21(DE3) Rosetta2 carrying pETGB1_HMGA1a were grown in M9 minimal media supplemented with, trace metals, thiamine, 0.5x MEM vitamin mix and ^15^N NH_4_ (1g/L) or ^15^N NH_4_ and ^13^C glucose. Protein expression was induced with 1.5mM IPTG and expressed at 20°C for 24h. The harvested cells were washed in RBS buffer (10mM HEPES pH 7.4, 10mM NaCl, 3mM MgCl2, 2mM PMSF) before storage at -80°C.

HIS6-GB1 tag of ^15^N or ^15^N^13^C labelled HMGA1a was cleaved using TEV protease (in-house) and the tag removed by passing the cleaved solution over Ni-NTA beads in 20mM K_2_PO_4_/KH_2_PO_4_ pH 7.2, 10mM imidazole, and 2mM β-mercaptoethanol. Cleaved HMGA1a was further purified by heparin chromatography, followed by size exclusion chromatography in 20mM K_2_HPO_4_/KH_2_PO_4_ pH 7, 150mM NaCl and 0.5mM β-mercaptoethanol. Final proteins were dialyzed into NMR buffer (1: 20mM Na_2_HPO_4_/NaH_2_PO_4_ pH 6.2, 0.1mM EDTA, 10 % D_2_O or 2: 20mM Na_2_HPO_4_/NaH_2_PO_4_ pH 6.2, 50 mM NaCl, 0.1mM EDTA, 10 % D_2_O).

RNAs were transcribed from annealed DNA oligonucleotides (8μM) using in-house T7 RNA polymerase in transcription buffer (40mM Tris-Cl pH8.1, 1mM Spermidine, 0.01% Triton X-100, 5mM DTT) supplemented with optimized concentration of MgCl_2_ (30-40 mM) for 3h. Reactions were stopped by the addition of EDTA. Unincorporated rNTPs were removed using G-25 PD1-columns (GE BioSciences), and full-length RNA was purified using HiTrap Q columns in 20mM Na_2_HPO_4_/NaH_2_PO_4_ pH 7.1, 0.1mM EDTA, 100-1000 mM NaCl. Purified fractions were pooled and dialyzed into 1mM K_2_HPO_4_/KH_2_PO_4_ 0.1mM EDTA. RNAs were diluted to 10-20 μM, heated to 65°C for 5min, flash folded in Liquid N_2_ and lyophilized. Final samples were resuspended in NMR buffer + 10 % D_2_O for analysis.

RNA sequences used in NMR:

SL2: GGCGUCCCUCCCGAAGCUGCGCGCUCGGUCGAAGAGGACGCC AU-rich dsRNA (PRDII): GGGAAATTCCTCTTCGGAGGAATTTCCC

NMR spectra were acquired on a Bruker AVII-600 MHz equipped with a cryoprobe. Data were processed on Topspin 4.0 (Bruker) and analyzed in NMRFAM-SPARKY ^72^. HMGA1a proteins were analyzed with ^1^H-^15^N-HSQC spectra. The folding of each RNA was analyzed by ^1^H ^1^H TOCSY at 25°C and ^1^H ^1^D spectra at 10°C optimized to detect protected imino resonances representing stable base pairs. To investigate the interaction between HMGA1a and RNA, RNA was titrated into HMGA1a protein until binding was saturated. Each step was monitored by ^1^H-^15^N-HSQC detecting ^1^H-^15^N resonances from HMGA1a and ^1^H ^1^D NMR spectra monitoring imino resonances between 10-16ppm. ^1^H-^15^N-HSQC of free HMGA1a (mouse) were compared to the human equivalent (Biological Magnetic Resonance Bank (BMRB) accession: 7279 ^73^, which showed excellent agreement enabling assignment of most residues except V75 (T in human) and A78 (T in human). The ^1^H-^15^N resonances were followed during the titrations to assign resonances of the RNA-bound forms. Assignments of both free HMGA1a and bound to 7SK SL2 RNA were confirmed with HNCACB, CBCACONH, HNCO. The amide backbone dynamics of HMGA1a bound to 7SK SL2 was further analyzed with interleaved ^15^N{^1^H}heteronuclear NOE. Combined chemical shift perturbations were calculated using the formula: CSP=[ΔH+(ΔN/7)/2].

## Interactome capture and RNP purification

UV-crosslinked HEK293T cells expressing HA-HMGA1 isoforms or HA-HMGA1a mutants were resuspended in lysis buffer (100mM Tris HCl, pH 7.5, 500mM LiCl, 10mM EDTA, pH 8.0, 1% lithium dodecyl sulphate, 5mM DTT) supplemented with Protease Inhibitor Cocktail (Roche). Cell lysis was accomplished by passing cell suspension through a 25-gauge needle. The cleared supernatant was incubated with Oligo d(T)_25_ beads at room temperature for 2h on a rotating wheel. The beads were washed twice with the lysis buffer at room temperature for 30min, then washed three times with wash buffer (50mM Tris HCl, pH 7.5, 140mM LiCl, 2mM EDTA, pH 8.0, 0.5% Igepal, 0.5mM DTT) at room temperature for 20min. The protein-bound RNAs were eluted by adding low salt elution buffer (10mM Tris HCl, pH 7.5) to the beads and heating the mixture at 80°C for 30min. The RNA was digested with 100U RNase I (Ambion for 30min at 37°C and the eluents were analyzed by Western blotting using a mouse anti-HA antibody (HA.C5, Thermo Fisher Scientific).

Covalently crosslinked protein-RNA complexes (XRNAX) were extracted from TRIzol® lysates as described in ^74^ (https://www.xrnax.com/theprotocol).

## Western blot

Whole cell lysates were prepared in RIPA buffer (50mM Tris-HCl, pH 8; 150mM NaCl, 1% Igepal, 0.5% sodium deoxycholate, 0.1% SDS) supplemented with Protease inhibitors (Roche). Ten micrograms of protein lysate were run on an Invitrogen™ NuPAGE™ Bis-Tris Mini Protein Gel, 4–12% with MES SDS Running Buffer (Thermo Fischer Scientific). The gel was subsequently transferred onto nitrocellulose membrane HMGA1 proteins were probed using an anti-HMGA1 rabbit monoclonal primary antibody (Abcam, ab129153). GAPDH protein was used as loading control (14C10, #2118, Cell Signaling Technologies). Primary antibody incubations were followed by secondary Goat Anti-Rabbit IgG (H + L)-HRP Conjugate (BioRad), and protein bands were visualized using Amersham™ ECL Western Blotting Detection Reagent (Cytiva Life Sciences) with ChemiDoc Imaging System (BioRad).

## Co-immunoprecipitation

Whole cell pellets from P19 cells transfected with HA-HMGA1-T2A-GFP vectors were lysed in ice-cold NET-2 buffer (50mM Tris-Cl pH 7.4, 150mM NaCl, 0.05% Igepal) supplemented with protease inhibitors (Merck). The lysates were sonicated and treated with 4U TURBO DNase I (Thermo Fisher Scientific) for 3min at 37°C 1100rpm to solubilize chromatin. A half of the cleared lysate was treated with 200ng RNase A and the other half with 80U RNase inhibitor (Thermo Fisher Scientific) for 10min at 37°C. Input samples (2%) were collected from each aliquot and the remaining volume was split into two aliquots for immunoprecipitation with Dynabeads Protein G (Thermo Fisher Scientific) coupled with either 2µg anti-HA antibody (Invitrogen, HA-C5) or 2µg mouse IgG isotype control (Invitrogen). Following overnight incubation at 4°C with rotation, the beads were washed four times with NET-2 buffer rotating for 1min at 4°C. The proteins were eluted with NuPAGE LDS Sample buffer (Thermo Fisher Scientific) supplemented with 0.1M dithiothreitol (DTT) at 85°C for 5min. The samples were separated by NuPAGE 4-12% Bis-Tris gels and transferred to a nitrocellulose membrane. The blots were probed with anti-HMGA1 (Abcam, ab129153) and anti-U170K (Abcam ab83306) and imaged on a ChemiDoc imaging system (Bio-Rad Laboratories, Hercules, CA, USA).

## HMGA1 knockdown, sequencing and computational analysis

HMGA1 was depleted in P19 cells by RNAi (75pmol siRNA ID S67598 or a scrambled siRNA control, Ambion) using Lipofectamine RNAiMAX Transfection Reagent (Thermo Fisher Scientific). Total protein and RNA were extracted 48h post-transfection for Western blot or sequencing library preparation, respectively. RNA was isolated using Tri-Reagent (Thermo Fisher Scientific), RNA was quantified using a Qubit RNA BR assay kit (Thermo Fisher Scientific), and RNA integrity was analyzed using Agilent 2100 bioanalyzer. Library preparation and RNA sequencing were performed at the Medical Genomics Facility (Monash Health Translation Precinct). Libraries were generated using an Illumina Stranded Total RNA-Seq Ribo-Zero kit. Sequencing was performed using paired-end sequencing using Illumina NextSeq550 high output mode and v2.5 chemistry with Illumina protocol 15046563 v04 (60 bp paired-end reads, 40M read pairs per sample).

For gene expression analysis, transcript abundances in short-read HMGA1 knockdown data were quantified using Salmon ^65^ with genome annotations from GENCODE 48 and m39 for the human and mouse data, respectively. Gene abundances were obtained using tximport ^75^ and differentially expressed genes between control and knockdown samples were determined with DEseq2 ^76^. A gene was considered differentially expressed with an adjusted p-value <0.05 and an absolute log_2_ fold change >1.

For splicing analysis, the reads were aligned to either the human or mouse reference genome, GENCODE 48 and m39, using HISAT2 ^77^. Differential splicing control and knockdown samples was determined using MAJIQ ^78^. MAJIQ was supplied with the appropriate GENCODE genome annotation and MAJIQ build and dPSI were ran with default parameters. VOILA modulize was ran to obtain splicing event types. An event was considered differentially spliced with a primary E(dPSI) >0.1 and a secondary E(dPSI) >0.05.

