## Supplementary tables and figures for "dirCLIP profiles variant-specific RNA-protein interactions via nanopore long-read sequencing"

**Supplementary Table 1.** SRSF3 binding sites and regions determined by dRNA-seq dirCLIP signal and read terminal analysis using either untreated (-UV) or UV-treated (+UV) transcriptome controls.

**Supplementary Table 2.** SRSF3 binding sites and regions determined by dcDNA-seq dirCLIP mutation profiling using either untreated or UV-treated transcriptome controls and including deletions (nucleotide coverage ≥50 and a mutation rate difference ≥0.03).

**Supplementary Table 3.** HMGA1 binding sites and regions determined by dRNA-seq dirCLIP signal and read terminal analysis using an untreated *in vitro* polyadenylated transcriptome control. For Nanocompore results of U1 RNA variants only selection is included (U1_variants).

**Supplementary Table 4. Detailed HMGA1a binding sites in U1 snRNA variants.**
Statistical data for HMGA1a dRNA-seq dirCLIP sites in U1 snRNA transcript variants. Sites were identified using Nanocompore comparison of HMGA1a dirCLIP to *in vitro* polyadenylated transcriptome control.

*A. U1 snRNA variant characteristics*

| Transcript ID | Gene ID | Symbol | Length (nt) | Read Count | Sites Detected |
|---|---|---|---|---|---|
| ENSMUST00000240501.1 | ENSMUSG00000119476.1 | Rnu1a1 | 164 | 1,717 | 5 |
| ENSMUST00001239531.1 | ENSMUSG00000118876.1 | Rnu1b1 | 165 | 884 | 2 |
| ENSMUST00000240438.1 | ENSMUSG00000119030.1 | Rnu1b2 | 165 | 902 | 2 |
| ENSMUST00000240510.1 | ENSMUSG00000118677.1 | Rnu1b6 | 166 | 1,217 | 6 |

*B. Significant binding sites by variant*

*U1a1 (Rnu1a1)*

| Position | Reference K-mer | GMM p-value | Log Odds Ratio | Structural Context |
|---|---|---|---|---|
| 24 | CATGA | $7.9 \times 10^{-5}$ | −0.625 | Stem-loop 1 |
| 43 | TTCCC | $3.8 \times 10^{-2}$ | +0.361 | Loop region |
| 86 | CCCTG | $8.4 \times 10^{-3}$ | −0.475 | Stem-loop 3 |
| 134 | AGTGG | $1.9 \times 10^{-2}$ | −0.394 | 3' region |
| 147 | GTTCG | $6.9 \times 10^{-3}$ | −0.867 | 3' region |

*U1b1 (Rnu1b1)*

| Position | Reference K-mer | GMM p-value | Log Odds Ratio | Structural Context |
|---|---|---|---|---|
| 24 | CATGA | $1.4 \times 10^{-3}$ | +0.747 | Stem-loop 1 |
| 89 | CTGCG | $3.7 \times 10^{-2}$ | +0.497 | Stem-loop 3 |

*U1b2 (Rnu1b2)*

| Position | Reference K-mer | GMM p-value | Log Odds Ratio | Structural Context |
|---|---|---|---|---|
| 24 | CATGA | $1.5 \times 10^{-3}$ | +0.710 | Stem-loop 1 |
| 89 | CTGCG | $9.7 \times 10^{-3}$ | +0.559 | Stem-loop 3 |

*U1b6 (Rnu1b6)*

| Position | Reference K-mer | GMM p-value | Log Odds Ratio | Structural Context |
|---|---|---|---|---|
| 19 | GATAC | $4.3 \times 10^{-2}$ | −0.535 | 5' region |
| 24 | CATGA | $1.4 \times 10^{-4}$ | −0.631 | Stem-loop 1 |
| 43 | TTCCC | $2.9 \times 10^{-2}$ | +1.070 | Loop region |
| 69 | ACTTT | $1.6 \times 10^{-2}$ | +0.468 | Internal region |
| 87 | CCCTG | $3.0 \times 10^{-2}$ | +0.457 | Stem-loop 3 |
| 89 | CTGCG | $3.7 \times 10^{-2}$ | +0.408 | Stem-loop 3 |

*C. Binding site conservation*

| Binding Pattern | Positions | Structural Location | Variants |
|---|---|---|---|
| Conserved (all variants) | 24, 86–89 | SL1, SL3 | U1a1, U1b1, U1b2, U1b6 |
| Partial (2 variants) | 43 | Loop | U1a1, U1b6 |
| U1a1-specific | 134, 147 | 3' region | U1a1 only |
| U1b6-specific | 19, 69 | 5' and internal | U1b6 only |

**Supplementary Table 5.** HMGA1 binding sites and regions determined by dcDNA-seq dirCLIP mutation profiling using either untreated or UV-treated transcriptome controls and including deletions (nucleotide coverage ≥50 and a mutation rate difference ≥0.03).

**Supplementary Table 6.** Differentially expressed genes following HMGA1 knockdown in murine P19 cells (DESeq2).

**Supplementary Table 7.** Splicing changes following HMGA1 knockdown in murine P19 cells (MAJIQ).

**Supplementary Table 8.** Differentially expressed genes following HMGA1 knockdown in human K562 cells/ENCODE (DESeq2).

**Supplementary Table 9.** Splicing changes following HMGA1 knockdown in human K562 cells/ENCODE (MAJIQ).

**Supplementary Table 10. Comparison of dirCLIP with established iCLIP and eCLIP methods.**

| Feature | iCLIP | eCLIP | dirCLIP (dRNA) | dirCLIP (dcDNA) |
|---|---|---|---|---|
| **Technical specifications** | | | | |
| Sequencing platform | Illumina | Illumina | ONT | ONT |
| Read length | 50–150 bp | 50–150 bp | Full-length | Full-length |
| PCR amplification | Yes | Yes | No | No |
| RNase fragmentation | Required | Required | Not required | Not required |
| Typical yield (reads) | 10–50M | 10–50M | 0.5–2M | 1–5M |
| Input RNA | 1–10 µg | 1–10 µg | 0.5–5 µg | 0.5–5 µg |
| **Detection capabilities** | | | | |
| Detection method | RT termination | RT termination + mutation | Signal + termination | Mutation profiling |
| Binding site resolution | Single nt | Single nt | ~5 nt | Single nt |
| Transcript coverage | 3' biased | 3' biased | 3' biased | Full-length |
| Isoform resolution | No | No | **Yes** | **Yes** |
| Native RNA modifications | Lost | Lost | **Preserved** | Lost |
| **Practical considerations** | | | | |
| Multiplexing | Yes | Yes | Limited | Limited |
| Cost per sample | $$ | $$ | $$$ | $$$ |
| Analysis pipelines | Established | Established | Custom | Custom |
| **Summary** | | | | |
| Key advantage | High throughput | Low background | Native RNA context | Full coverage |
| Key limitation | No isoform context | No isoform context | 5' coverage bias | RT-dependent |

ONT, Oxford Nanopore Technologies; RT, reverse transcriptase; nt, nucleotide. iCLIP, individual-nucleotide resolution CLIP [1]; eCLIP, enhanced CLIP [2]. Cost estimates: $$, $200–500 per sample; $$$, $500–1000 per sample (consumables only, excluding equipment). Read yields are approximate and depend on experimental conditions and sequencing depth.

### SUPPLEMENTARY FIGURES AND LEGENDS

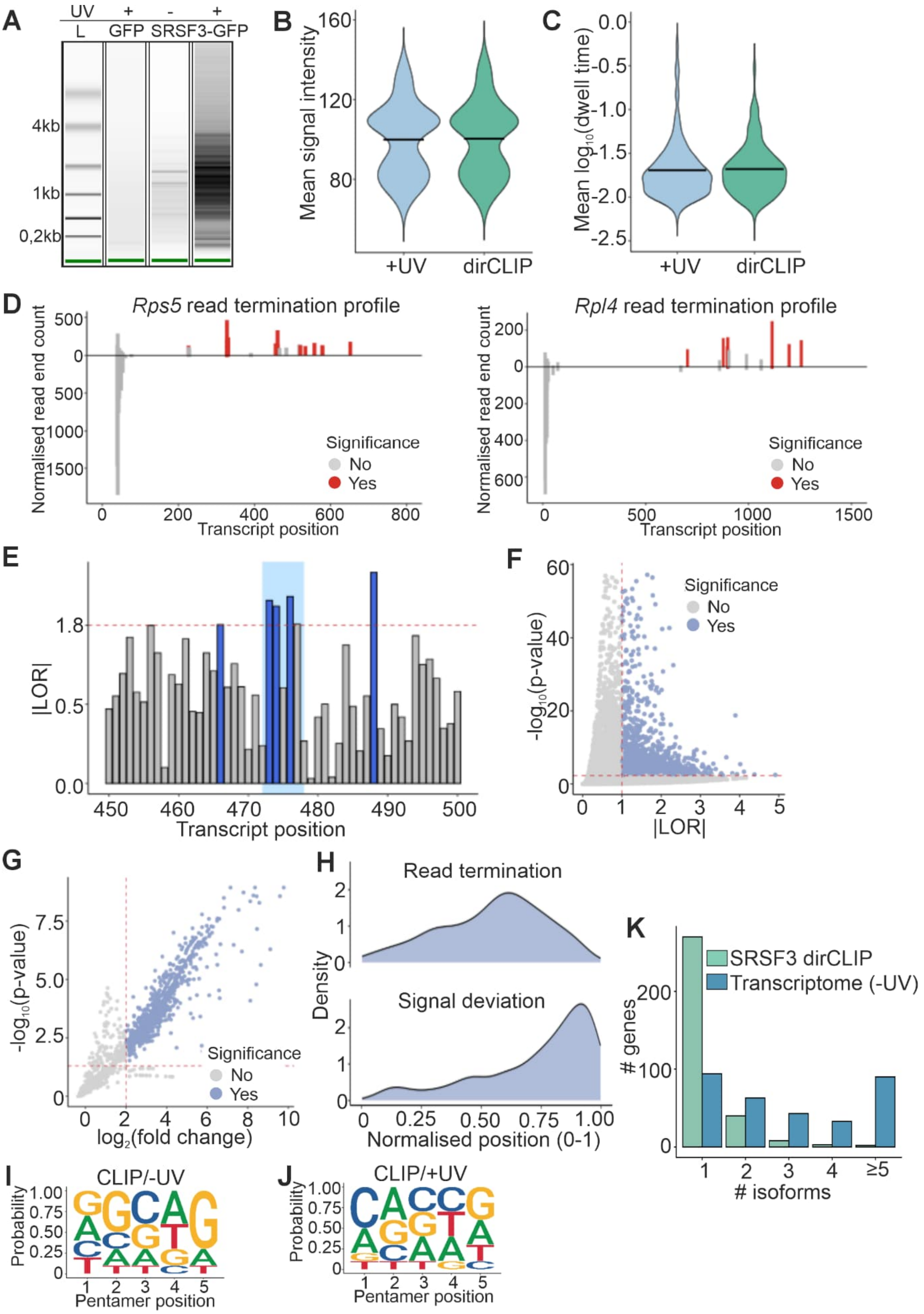


**Supplementary Figure 1. Details of direct RNA sequencing-based detection of SRSF3**

**RNA binding sites.** (**A**) UV-dependent immunoprecipitation of SRSF3-bound long RNAs in P19 cells expressing SRSF3-GFP or nuclear GFP as a control. L=ladder. **(B)** Mean signal intensity of significant sites detected with Nanocompore comparison of SRSF3 and UV-treated transcriptome (+UV). Horizontal bars indicate mean values per condition. (**C**) Same as (A), but for dwell time. (**D**) Library size-normalized read end counts per million (CPM) for select representative transcripts *Rps5* and *Rpl4*. SRSF3 dirCLIP counts are plotted above the axis and UV-treated transcriptome counts below the axis, with significant differential sites marked in red. (**E**) Excerpt of absolute Log Odds Ratios (|LOR) for transcript *Rps13* at positions 450-500nt showing several significant (light blue) sites clustered close together, likely due to a single crosslinking event. **(F)** Significant differences in signal intensity and dwell time (signal deviation) between SRSF3 dirCLIP and untreated (-UV) transcriptome samples identified by Nanocompore. (**G**) Significant differences in read end termination rates between SRSF3 and untreated (-UV) transcriptome samples detected by Poisson GLM models. (**H**) Density distribution of significant sites identified through signal deviation and read termination methods across normalized transcript coordinates when using untreated (-UV) transcriptome control. (**I**) Sequence logo plot of enriched motifs (z-score >2 and p<0.05) derived from genomic ranges of ±50 bp around the significant SRSF3 dirCLIP/untreated transcriptome (-UV) sites. (**J**) Same as (I), but for CLIP/UV-treated transcriptome (+UV) sites. **(K)** Comparison of the number of isoforms per gene with significant SRSF3 binding sites derived from SRSF3 dirCLIP versus the number of expressed isoforms (TPM > 0.5) for those same genes in the untreated (-UV) transcriptome sample.

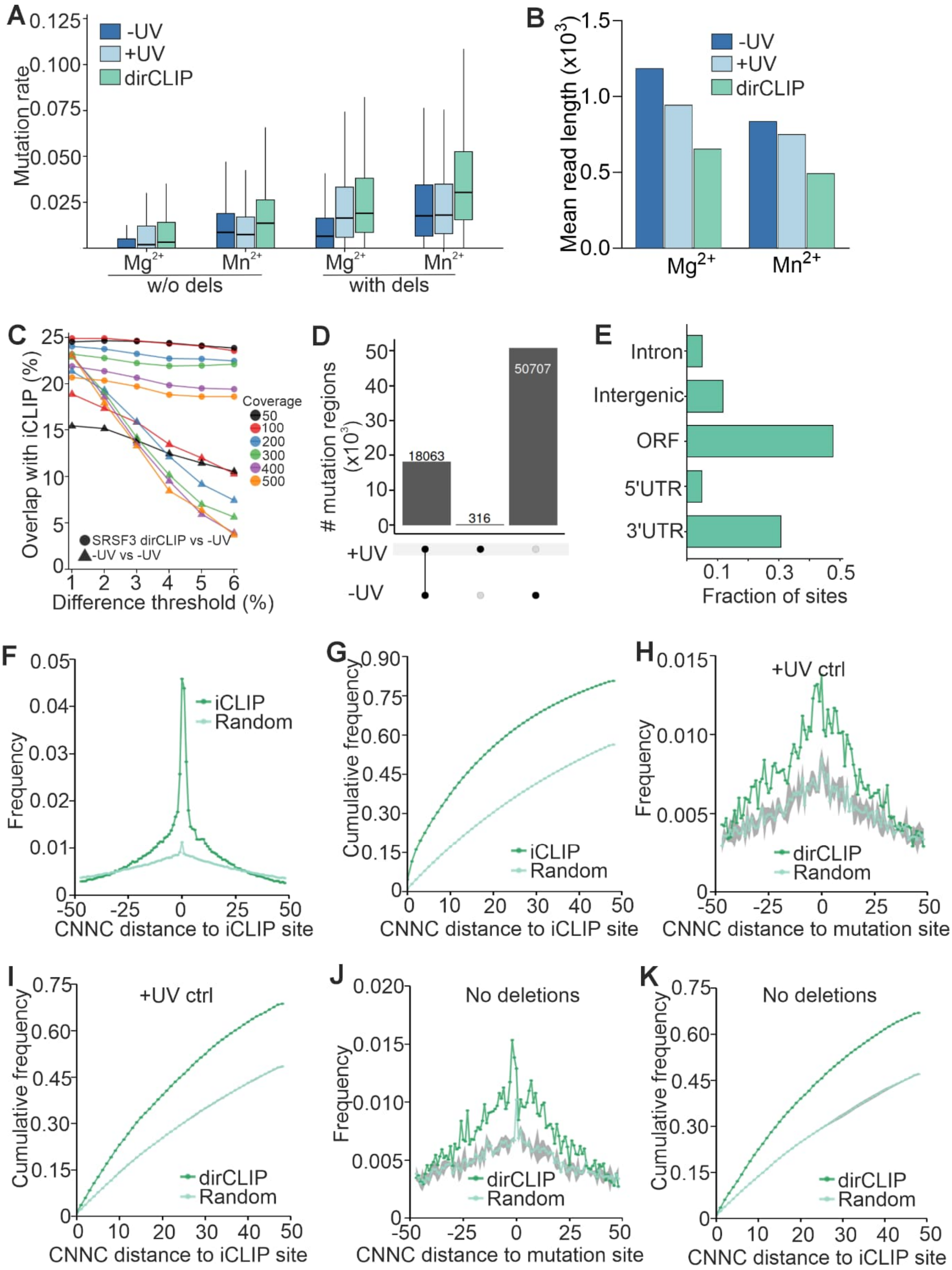


**Supplementary Figure 2. Details of direct cDNA sequencing-based detection of SRSF3 RNA binding sites.** (**A**) dcDNA-seq mutation rate per nucleotide with and without (w/o) deletions (dels) for untreated transcriptome (-UV), UV-treated transcriptome (+UV), and

SRSF3 dirCLIP samples in $MnCl_2$ buffer conditions. Nucleotides with a coverage ≥50 were included. (**B**) Mean read length of untreated (-UV), UV-treated (+UV) dcDNA-seq control transcriptomes and dirCLIP samples prepared in $MgCl_2$ and $MnCl_2$ buffer conditions. Primary alignment reads >20 nucleotides or <99th percentile were used. (**C**) The percentage of significant SRSF3 dcDNA-seq dirCLIP sites that overlap with SRSF3 iCLIP peaks identified using the iCount. Untreated transcriptome was used as the control and deletions were included in the analysis. The x-axis indicates mutation rate difference thresholds. The solid circles represent data points for SRSF3 dirCLIP using the untreated transcriptome as a control and the solid triangles (-UV) untreated transcriptome to transcriptome comparison (random sites). (**D**) Significant SRSF3 dcDNA-seq dirCLIP regions identified using untreated transcriptome (-UV) or UV-treated transcriptome (+UV) controls. Mutation rates were calculated including deletions and significant sites defined using a nucleotide coverage ≥50 and a mutation rate difference ≥0.03. Significant mutation sites located within five base pairs of each other were merged into a single continuous region. (**E**) Distribution of significant SRSF3 dcDNA-seq dirCLIP sites across genome regions. Significant sites were defined as in Figure 2. (**F**) The frequency distribution of SRSF3 iCLIP peak center distance to the closest CNNC motif (top ten motifs with highest z-scores identified from SRSF3 iCLIP peaks). Light green line and scatter plot represent the same analysis using sequences in which the ±50 bp window around each peak center was randomly shuffled. The gray shaded area indicates the standard error of the mean from hundred rounds of random shuffling. (**G**) Cumulative frequency distribution of SRSF3 iCLIP peak center distance to the closest CNNC motif as in (F). (**H**) Distance between SRSF3 dcDNA-seq dirCLIP sites and CNNC motifs. As a control, sequences within ±50nt window around each dirCLIP site were randomly shuffled (random). The gray shaded area indicates the standard error of the mean from 100 rounds of random shuffling. Significant dirCLIP sites were identified using the UV-treated transcriptome (+UV) as a control with a nucleotide coverage ≥50 and a mutation rate difference ≥0.03 including deletions in the mutation rate calculation. (**I**) Cumulative frequency distribution of SRSF3 dcDNA-seq dirCLIP site distance to the closest CNNC motif as in (H). (**J**) Distance between SRSF3 dcDNA-seq dirCLIP sites and CNNC motifs as in (H) when significant dirCLIP sites were identified using the untreated transcriptome as a control excluding deletions in the mutation rate calculation. (**K**) Cumulative frequency distribution of SRSF3 dcDNA-seq dirCLIP site distance to the closest CNNC motif as in (J).

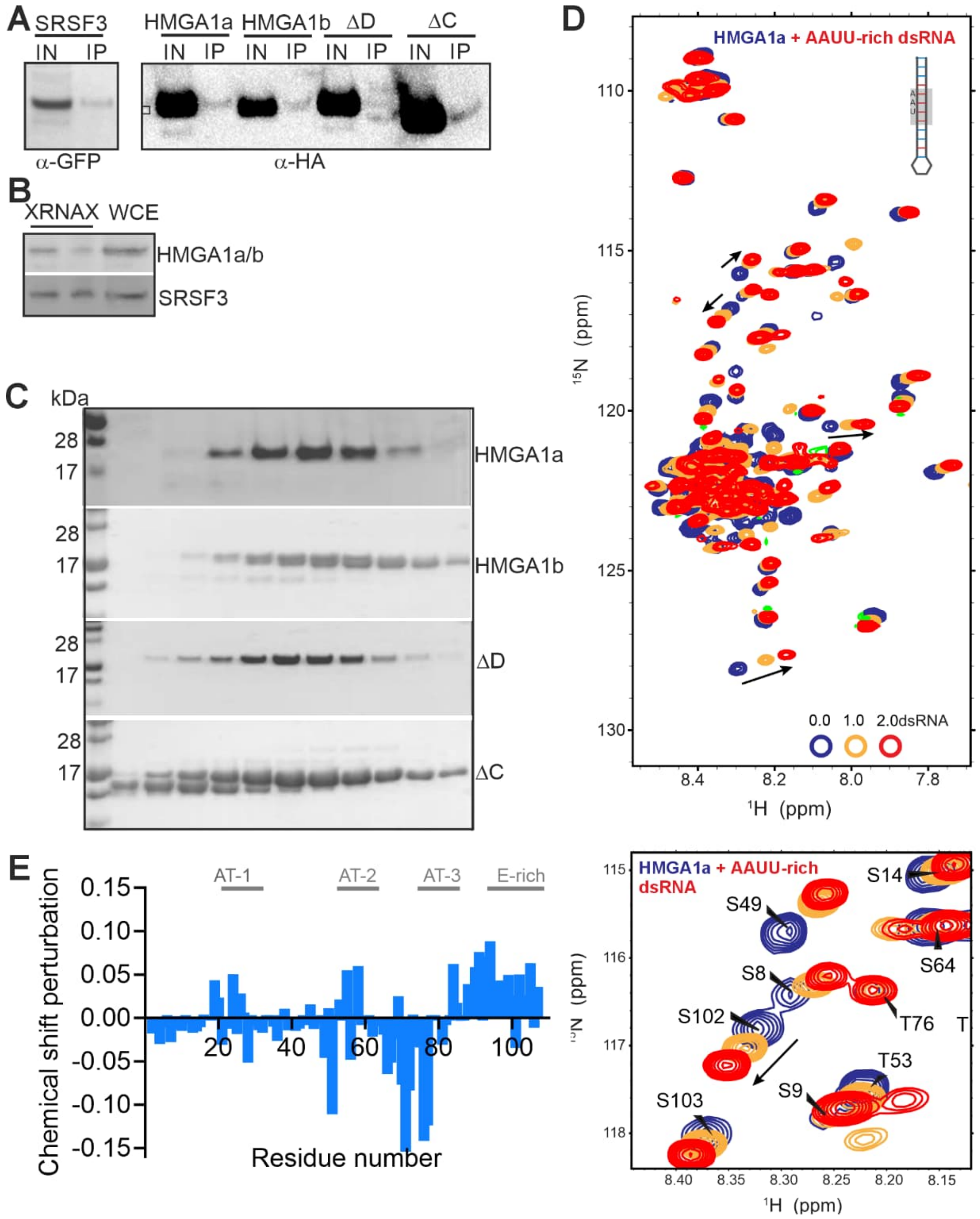


**Supplementary Figure 3. HMGA1a and b bind to RNA via AT-hooks.** (**A**) Interactome capture of poly(A) mRNA in HEK293 cells expressing HA-tagged HMGA1 isoforms a or b, or HMGA1a DNA binding (ΔD) and C-terminal deletion (ΔC) mutants. Interactome capture of poly(A) mRNA in cells expressing SRSF3-EGFP shown as positive control. (**B**) Western blot of ribonucleoprotein complexes purified from P19 cells using XRNAX. WCE, whole cell extract. (**C**) SDS-PAGE of purified HMGA1 isoforms a and b, and HMGA1a DNA binding (ΔD) and C-terminal deletion (ΔC) mutants. (**D**) $^{1}H$-$^{15}N$ HSQC NMR spectra of free HMGA1a (blue) and HMGA1a bound to an AAUU-rich dsRNA derived from the PRDII sequence of the INF-β promoter (yellow, molar equivalent; red, 2.0 molar equivalent). The black arrows indicate the chemical shift changes observed when RNA was added, demonstrating RNA interaction. Enlarged sections of the spectra below display chemical shift changes in residues S102 and S49. (**E**) Combined chemical shift perturbation plots of N-H signals obtained from the $^{1}H$-$^{15}N$ HSQC NMR spectra in (D).

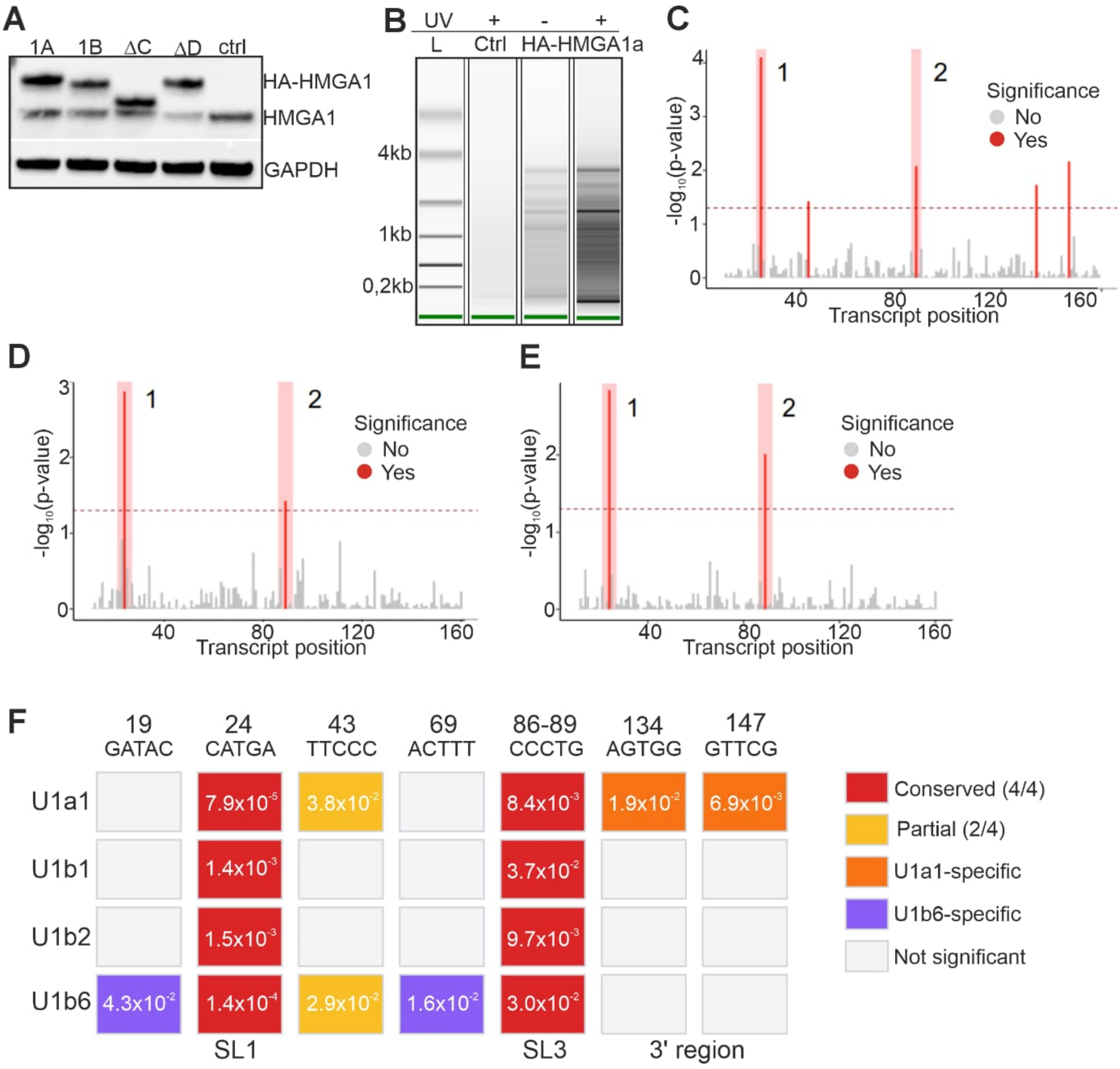


**Supplementary Figure 4. Details of dRNA-seq dirCLIP identification of HMGA1 RNA binding sites.** (**A**) Expression of HA-tagged HMGA1 isoforms a or b, or HMGA1a DNA binding (ΔD) and C-terminal deletion (ΔC) mutants in P19 cells. (**B**) UV-dependent immunoprecipitation of HMGA1a-bound long RNAs in P19 cells expressing HA-tagged HMGA1a or control. L=ladder. Distribution of dRNA-seq dirCLIP sites meeting significance threshold in (**C**) U1a1, (**D**) U1b1 and (**E**) U1b2 snRNA variants. Sites marked in pink were detected in multiple U1 snRNA variants. (**F**) Conservation matrix showing binding site significance across variants. Cell colors indicate conservation pattern; numbers indicate GMM p-values (significant sites GMM p-value<0.05). Grey cells indicate non-significant sites. SL, stem-loop.

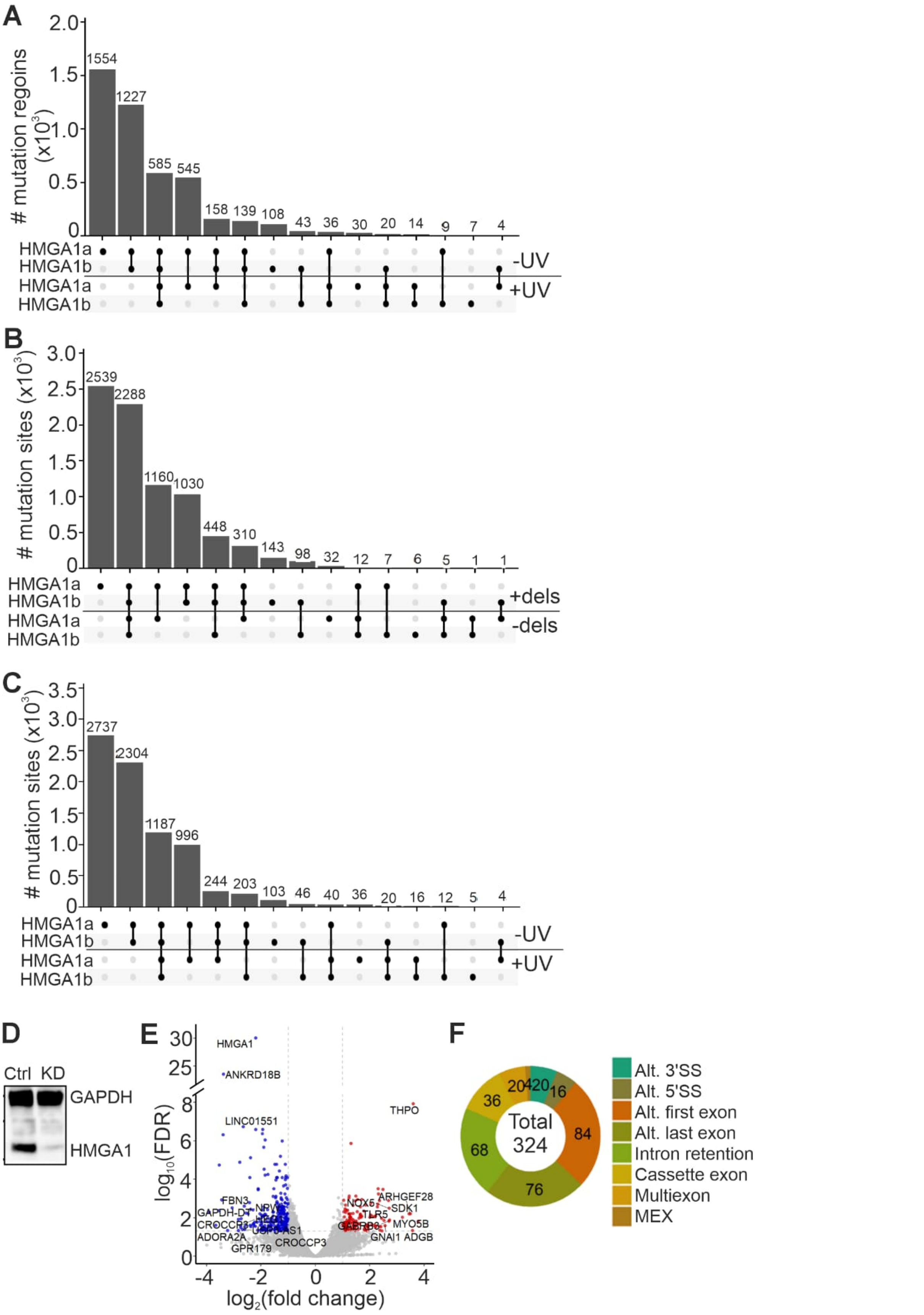


**Supplementary Figure 5. Details of dcDNA-seq dirCLIP identification of HMGA1 RNA binding sites.** (**A**) Significant dcDNA dirCLIP regions in HMGA1a and b dirCLIP samples identified using untreated transcriptome (-UV) or UV-treated transcriptome (+UV) controls. Mutation rates were calculated including deletions and significant sites defined using a nucleotide coverage ≥50 and a mutation rate ≥ 0.03. Significant mutation sites located within five base pairs of each other were merged into a single continuous region. (**B**) Significant dcDNA dirCLIP regions in HMGA1a and b dirCLIP samples identified using untreated transcriptome (-UV) controls including (+dels) or excluding (-dels) deletions in the mutation rate calculations. Significant sites were defined using a nucleotide coverage ≥50 and a mutation rate difference ≥0.03. (**C**) Coverage-normalized significant dcDNA-seq dirCLIP sites for HMGA1a and HMGA1b, demonstrating higher RNA binding activity of the HMGA1a isoform. (**D**) Western blot of P19 cells treated with HMGA1 siRNA or a scrambled control. (**E**) Differentially expressed genes following HMGA1 CRISPRi in human K562 cells (ENCODE project: ENCSR069OUL). (**F**) Splicing changes following HMGA1 CRISPRi in human K562 cells (ENCODE project: ENCSR069OUL).
